# Annotation of Chromatin States in 66 Complete Mouse Epigenomes During Development

**DOI:** 10.1101/2020.07.23.218552

**Authors:** Arjan van der Velde, Kaili Fan, Junko Tsuji, Jill Moore, Michael Purcaro, Henry Pratt, Zhiping Weng

## Abstract

The morphologically and functionally distinct cell types of a multicellular organism are maintained by epigenomes and gene expression programs. Phase III of the ENCODE Project profiled 66 mouse epigenomes across twelve tissues at daily intervals from embryonic day 10.5 to birth. Applying the ChromHMM algorithm to these epigenomes, we annotated eighteen chromatin states with characteristics of promoters, enhancers, transcribed regions, repressed regions, and quiescent regions throughout the developmental time course. Our integrative analyses delineate the tissue specificity and developmental trajectory of the loci in these chromatin states. Approximately 0.3% of each epigenome is assigned to a bivalent chromatin state, which harbors both active marks and the repressive mark H3K27me3. Highly evolutionarily conserved, these loci are enriched in silencers bound by Polycomb Repressive Complex proteins and the transcription start sites of their silenced target genes. This collection of chromatin state assignments provides a useful resource for studying mammalian development.

## INTRODUCTION

Multicellular organisms maintain a myriad of cell types along separate lineages to carry out the cellular programs required for development and survival. These cell types all have the same genome but different epigenomes, characterized by chromatin accessibility, histone modifications, and DNA methylation, which cooperate with trans-factors to regulate gene expression and downstream activities. Thus, systematic annotation of epigenomes is essential for understanding genome functions. Experimental techniques such as chromatin immunoprecipitation followed by sequencing (ChIP-seq)^1–3^, transposase accessible chromatin with sequencing (ATAC-seq)^4^, and whole-genome bisulfite sequencing (WGBS)^5^ enable genome-wide profiling of histone marks, chromatin accessibility, and DNA methylation, respectively. When several of these epigenetic marks have been profiled for a given cell type, the results can be integrated using computational algorithms such as ChromHMM^6^, Segway^7^, and IDEAS^8^ to classify genomic loci into a small number of chromatin states, such that the chromatin state of a locus is predictive of its function in the given cell type.

Coordinated efforts by the ENCODE and Roadmap Epigenomics Consortia provided tremendous insights into gene regulation in a diverse array of human cell and tissue types^9,10^. The mouseENCODE project furthered our understanding of multiple adult mouse cell types^11^. Phase III of the ENCODE Consortium generated 66 complete mouse epigenomes across 12 fetal tissues at four to seven developmental time-points with a daily interval, each investigated with ten assays^12^: ATAC-seq^13^, WGBS^14^, and ChIP-seq of eight histone marks^13^. The histone marks included histone 3 lysine 4 trimethylation (H3K4me3) and histone 3 lysine 9 acetylation (H3K9ac), enriched at promoters and present at enhancers^1,15–17^; H3K27ac, H3K4me1, and H3K4me2, enriched at enhancers^1,15,17,18^; H3K36me3, enriched within the bodies of actively transcribed genes^19^; H3K27me3, catalyzed by and guiding the Polycomb Repressive Complexes (PRC) of proteins to repress gene expression^20^; and H3K9me3, enriched in heterochromatin to silence repeats and gene clusters^19^. All these 66 epigenomes were accompanied by transcriptome sequencing (RNA-seq) data^21^, and 20 of the biosamples were assayed by DNase-seq, another technique for measuring chromatin accessibility^22^ (**Fig. 1a** and **Supplementary Table 1**). This body of data was generated by four ENCODE labs, with the same type of data generated by the same lab, representing the most complete epigenetic data on fetal mouse tissues, ideal for characterizing the epigenomic landscape during mammalian development.

**Figure 1:**
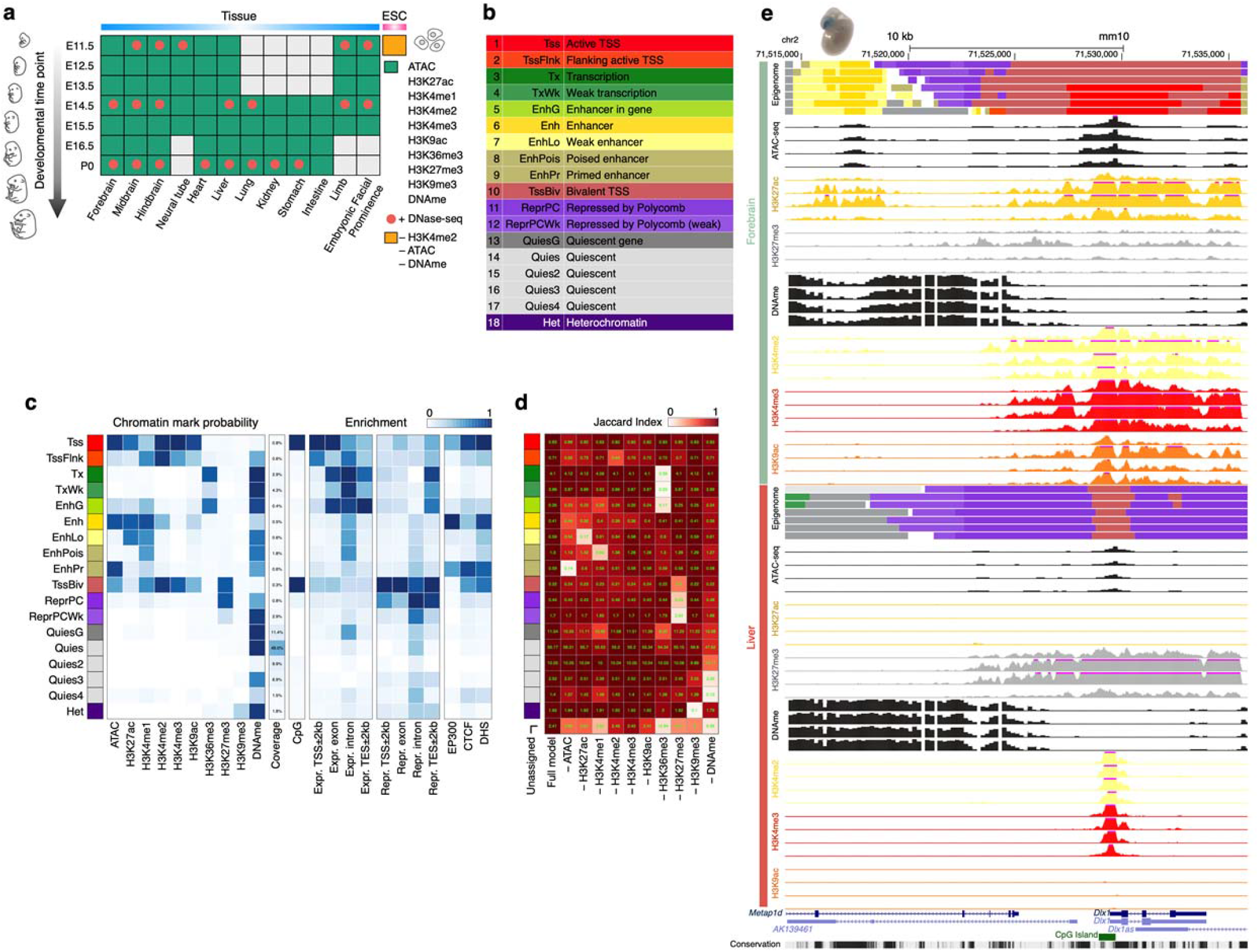
Overview of the 66 epigenomes and 18 chromatin states during mouse embryogenesis. **a.** Twelve tissues at 4-7 developmental time-points have ChIP-seq data for eight histone marks (green boxes), ATAC-seq data, and DNA methylation (DNAme) data, totaling 66 complete epigenomes. Twenty-one of these epigenomes also have DNase-seq data (red dots). Embryonic stem cells (orange box) have ChIP-seq data for seven histone marks, and are missing H3K4me2, ATAC-seq, and DNAme. **b.** Eighteen chromatin states are defined by ChromHMM across the 66 complete epigenomes. **c.** Histone-mark probabilities, genome coverage, and overlapping genomic features including gene expression, regulatory features (P300 binding, CTCF binding, and DNase I hypersensitive sites), and distances to the TSSs of expressed and repressed genes are shown for each chromatin state. The enrichments for the categories are the averaged values across tissues and time-points. **d.** Jaccard similarities between the partial epigenomes with each mark omitted and the ten-mark E13.5 midbrain epigenome. **e.** The *Dlx1* locus is displayed with chromatin states (color-coded as in **a**) in the forebrain and the liver for all seven time-points. Also shown are the signals of several histone marks (scale: 0-50) that differ between forebrain and liver (for E11.5, E13.5, E15.5, and P0 only, due to space constraints), along with ATAC and DNA methylation signals. A transgenic mouse embryo is shown on top of the enhancer region, indicating the forebrain-specific activity of this enhancer. A CpG island that overlaps with the bivalent region at the TSS of *Dlx1* is shown at the bottom of the panel.

We applied ChromHMM^6^ to these 66 mouse epigenomes and defined 18 chromatin states (**Fig. 1b**). Most of these mouse chromatin states recapitulated the 15 human chromatin states defined by the Roadmap Epigenomics Consortium using a subset of five histone marks in human biosamples^10^, and our novel states corresponded to a refinement of previously defined enhancer, bivalent, and quiescent states. We observed a substantially larger variation of chromatin state assignments among the mouse tissue types at a given developmental time-point than we did across all developmental time-points for a single tissue. We further investigated one chromatin state in detail—TssBiv, a bivalent state enriched in the transcription start sites (TSS) which harbors both active marks (H3K4me3, H3K4me2, H3K4me1, and H3K9ac) and the repressive mark H3K27me3. We found that genomic loci in TssBiv were substantially more evolutionarily conserved than loci in any of the other 17 Chromatin states. Genes with bivalent TSSs were first identified in embryonic stem cells and thought to be poised for activation or repression in response to developmental or environmental cues^23^. Subsequently, such bivalent domains were reported in differentiated cell types^24–27^, but they have not been studied during fetal development. Each fetal tissue harbors approximately 3000 bivalent genes and they are repressed in expression in that specific tissue. These bivalent genes are highly enriched in transcription factors (TFs) differentially expressed among the fetal tissues. Comparison with recently defined silencers bound by the Polycomb Repressive Complex 2 (PRC2) proteins^28^ revealed that both the PRC2-bound silencers and the TSSs of their silenced genes are highly enriched in the bivalent regions. Thus, the bivalent regions support an evolutionarily conserved silencing mechanism for lineage-specific genes, in particular the master TFs controlling tissue development. Our comprehensive annotation of chromatin states provides a resource for studying mammalian development.

## RESULTS

### Chromatin states were defined using ATAC-seq, WGBS, and the ChIP-seq data of eight histone marks

The 66 mouse fetal epigenomes, all complete with ten chromatin marks, represent a comprehensive collection for chromatin state assignment (**Fig. 1a**). We used ChromHMM to learn 18 states jointly from this dataset (**Fig. 1b, c**). ChromHMM chunks the genome into non-overlapping 200 base-pair (bp) bins and assigns each of these genomic bins to one of the 18 chromatin states in each biosample. We named our chromatin states in a way to be consistent with earlier ChromHMM publications^6,10,29^. Two of our learned states are proximal to active TSSs (Tss and TssFlnk, approximately 1.5% of the mouse genome); two states associate with actively transcribed genes (Tx and TxWk, 8.5%); five states are enhancer-related (Enh, EnhLo, EnhPois, EnhPr, and EnhG; 4.5%); one bivalent state often falls near inactive TSSs (TssBiv, 0.3%); three states are repressive (ReprPC and ReprPCWk enriched in H3K27me3, 5.5%; and Het in H3K9me3, 2.5%); and five states are quiescent (QuiesG, Quies, Quies2, Quies3, and Quies4; 75%). The remaining 2% or so of the genome could not be confidently assigned to any one state.

The assignments of the 18 states are supported by comparison with gene expression and epigenomic data available for a subset of biosamples (**Supplementary Table 1**). Although both the active-TSS (Tss) and the bivalent-TSS states (TssBiv) are highly enriched in CpG islands, Tss (along with the TSS-flanking state TssFlnk) is only enriched in the TSSs of expressed genes (determined using RNA-seq data in the corresponding biosample) while TssBiv is only enriched in the TSSs of the repressed genes (**Fig. 1c**). The transcription-related states (Tx and TxWk) are enriched in the exons and introns of expressed genes but not those of the repressed genes (**Fig. 1c**). Enh (high-signal enhancer) is the state most enriched in the ChIP-signal of EP300, a histone acetyltransferase that preferentially binds active enhancers^30,31^ (**Fig. 1c**). The relative enrichment of the 18 states for the ATAC signal are highly consistent with their enrichments in DNase hypersensitive sites (DHS), determined using DNase-seq data in the corresponding biosample (**Fig. 1c**).

### Contributions of the chromatin marks to the assignments of chromatin states

To assess the contribution made by each of the eight histone marks, ATAC, and DNA methylation, we asked how accurately the ten-mark model would be able to annotate a new epigenome missing data for one of the marks. We addressed this question by removing the data for each mark individually from the midbrain E13.5 epigenome and computing the Jaccard similarity index between the chromatin state assignments of all genomic bins (each 200 bp long, which is the resolution of ChromHMM) with the data for the remaining nine marks. If a genomic bin has a posterior probability less than 0.5, then it is classified as unassigned. In general, when a mark is removed, the states most severely affected were among those states most enriched in this mark in the ten-mark model (compare **Fig. 1d** and the chromatin-mark probabilities in **1c**). However, the converse is not necessarily true, reflecting the redundancy between the marks. For example, the removal of H3K27ac affects the low-signal enhancer state (EnhLo) although the high-signal enhancer (Enh) state is even more enriched in H3K27ac than EnhLo (**Fig. 1c-d**). H3K4me3 and H3K9ac, when removed individually, did not have a major impact on any of the states although promoter states are enriched in H3K4me3 and both promoter and enhancer states are enriched in H3K9ac (**Fig. 1c-d**), indicating that the information contained by each of these two marks is already accounted for by the other nine marks. On the other hand, H3K36me3, H3K27me3, and H3K9me3 each brings non-redundant information to the ten-mark model, as all the states enriched in each of these marks were affected when the mark was removed (**Fig. 1c-d**).

### Chromatin states are conserved between human and mouse

The Roadmap Epigenomics Consortium previously defined 15 human chromatin states using five histone marks in 127 human biosamples^10^. To investigate the conservation of chromatin state types between human and mouse, we built a 15 state model using the same set of five histone marks in the 66 mouse fetal biosamples. This five-mark 15-state mouse model recapitulated 13 of the 15 human states identified by the Roadmap Epigenomics Consortium, with nearly identical emission probabilities and similar genome coverages. The 13 reproduced states, including the promoter, enhancer, transcribed, repressed, and bivalent states, were enriched in at least one of the five histone marks.

The remaining two mouse chromatin states had similar chromatin-mark probabilities to, but different genome coverages from, the two remaining human states^10^ (**Supplementary Fig. 1a**). These human states—the weak transcription state TxWk and the weak repressed polycomb state ReprPCWk (11.6% and 8.3% of the human genome)—had low signals for all five marks, and their assignments were based on their weak enrichments in expressed and repressed genes, respectively^10^. We identified a similar state with low signals for all marks in mouse, but although it was enriched within gene bodies in general, it was not enriched in either expressed or repressed genes in particular. We thus denoted it as the quiescent gene state (QuiesG, 25.17% of the mouse genome). We also identified a minor state (0.13% of the mouse genome) marked by both H3K36me3 and H3K27me3; we denote this state TxWk because regions in this state were assigned to the transcription state (Tx), the repressed polycomb state (ReprPC), or the weak repressed polycomb state (ReprPCWk) in our complete ten-mark, 18-state model. In summary, our results indicate that the chromatin states are highly conserved between human and mouse, and ChromHMM is able to identify these states reliably.

### Addition of three more histone marks, chromatin accessibility, and DNA methylation further clarified enhancer, bivalent, and quiescent states

To investigate the impact of incorporating additional data in the annotation of chromatin states, we constructed a 15-state model using all eight available histone marks (**Supplementary Fig. 1a**). We compared this model, and the five-mark 15-state model described above, with our 18-state model that further incorporated chromatin accessibility and DNA methylation data (ten marks in total, **Fig. 1**, also included in **Supplementary Fig. 1a** to facilitate comparison with the five-mark and eight-mark models). Comparison of the three ChromHMM models built with increasing numbers of epigenetic marks (five, eight, and ten marks) revealed that assignments differ predominantly for the enhancer, bivalent, and quiescent states (**Supplementary Fig. 1b-e**).

The five-mark model specified one enhancer state (Enh; 3.7% of the mouse genome) with high H3K4me1 levels (**Supplementary Fig. 1a**). Genomic regions in this state were assigned to five distinct enhancer states in the eight-mark model, which reflected different levels of three additional enhancer marks (H3K4me2, H3K9ac, and H3K27ac). Among these five states in the eight-mark model, the high-signal enhancer state Enh, which showed high levels for all these four enhancer marks, occupied only 0.2% of the genome (**Supplementary Fig. 1a, d**). The high-signal enhancer state Enh defined by the ten-mark model further showed high chromatin accessibility (ATAC signal) and low DNA methylation, occupying 0.64% of the genome (**Supplementary Fig. 1a, d**). The ten-mark model defined three additional enhancer states, with two of the three (EnhLo and EnhPois) being regroupings of the genomic regions assigned to the four enhancer states in the eight-mark model. The other enhancer state defined by the ten-mark model (EnhPr) corresponded to a subset of the regions assigned one of the enhancer states by the eight-mark model, showing high chromatin accessibility but low levels of enhancer marks (**Supplementary Fig. 1d**). Thus, the additional marks led to refined definitions of enhancer states.

One example of a tissue-specific enhancer is located inside the housekeeping gene *Metap1d* (methionyl aminopeptidase Type 1D, which functions in the mitochondria) and 10 kb upstream of the *Dlx1* gene, which encodes a brain-specific homeobox transcription factor. *Dlx1* is highly expressed in the forebrain (~200 transcripts per million or TPM), but not expressed in most other tissues (e.g., < 3 TPM in the liver). This region is annotated as a high-signal enhancer (Enh) in the forebrain, showing high ATAC and H3K27ac signals and low DNA methylation. It is annotated as a quiescent gene (QuiesG) in the liver due to its low ATAC and histone mark signals and high DNA methylation (**Fig. 1e**). A VISTA enhancer (accession: hs553) overlaps this region, and it is active in the forebrain and cranial nerve of mouse embryos^32^.

The five-mark model annotated three bivalent states with high levels of the active marks H3K4me1 and H3K4me3, as well as high levels of the repressive mark H3K27me3; however, both the eight-mark and ten-mark models only annotated one bivalent state, which additionally showed high levels of other active marks (H3K4me2, H3K9ac, and ATAC) and low levels of DNA methylation (**Supplementary Fig. 1a**). Roughly the same set of genomic regions were assigned to these bivalent states across the three models, suggesting that the state definition became more complete when more marks were available (**Supplementary Fig. 1e**).

The five-mark and eight-mark models annotated one quiescent state (Quies), which had very low signals for all available histone marks. The ten-mark model defined three additional quiescent states besides Quies. These four quiescent states all showed very low levels of the eight histone marks and ATAC, but they differed in DNA methylation, with the Quies state (49.0% of the genome) showing very high methylation levels and the Quies2 state (9.9% of the genome) showing very low levels of DNA methylation (**Supplementary Fig. 1a**). The quiescent states in the three models cover roughly the same set of genomic regions (**Supplementary Fig. 1b**).

### Variation of state assignments across tissues and along developmental time-points

After carefully analyzing the properties of the chromatin states in the ten-mark model, we assessed how variable the assignments of these states were among the 66 mouse epigenomes. We computed the Jaccard similarity index on the genomic regions assigned to each state between tissues or between developmental time-points. The enhancer states exhibited the greatest variability among tissues or across time-points, while the promoter, quiescent, and transcription states showed the least variability (**Fig. 2a**). The repressive state Het, enriched in H3K9me3, was almost as variable as the enhancer states (**Fig. 2a**). Moreover, all chromatin states were more similar across time-points in the same tissue than across tissues at the same time-point (**Fig. 2a**), consistent with the notion that the epigenome is inherited within the cell lineage.

**Figure 2:**
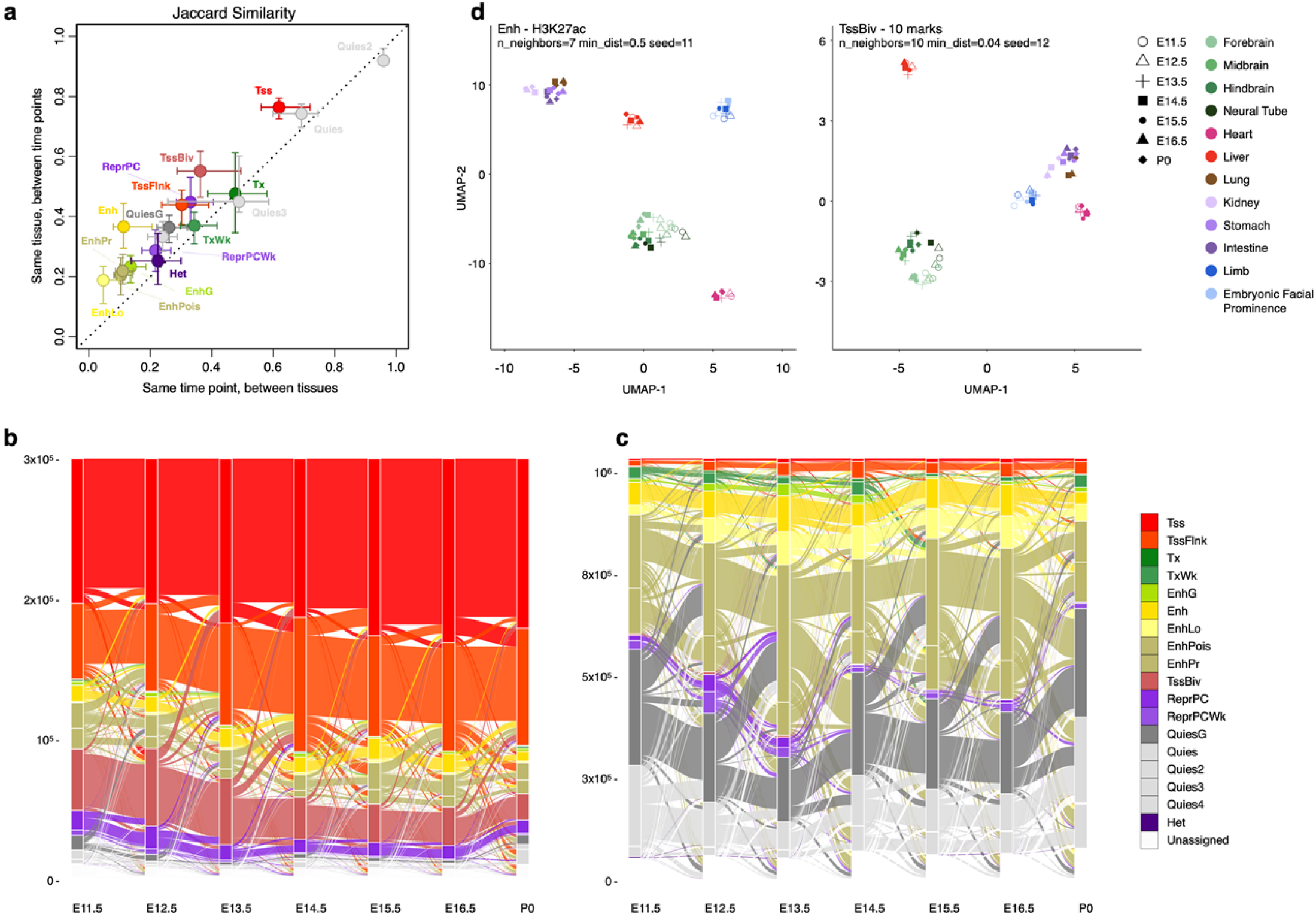
Variations of the chromatin states across tissues and their transitions along the developmental trajectory. **a.** Jaccard similarity between different time-points in the same tissue (y-axis) versus the similarity between different tissues at the same time-point (x-axis). Error bars indicate the range between the first and third quartiles. **b.** Transitions between chromatin states along midbrain developmental time-points. For clarity, only the genomic bins assigned TSS-related states (Tss, TssFlnk, and TssBiv) at one or more time-points are included. **c.** Same as b but for genomic bins assigned enhancer-related states (Enh, EnhLo, EnhPois, and EnhPr) at one or more developmental time-points. **d.** Visualization of the 66 epigenomes in two dimensions using the UMAP technique. (Left) UMAP was given the H3K27ac signals in the Enh genomic bins across the 66 epigenomes. There were 735,048 such genomic bins, which were assigned Enh in one or more epigenomes. (Right) UMAP was given the signals of all ten marks in the TssBiv genomic bins across the 66 epigenomes. There were 156,752 such genomic bins, which were assigned TssBiv in one or more epigenomes.

Temporal chromatin state transitions for each tissue occurred mostly between related states, e.g., among the promoter states (Tss, TssFlnk, and TssBiv) or among the enhancer states (Enh, EnhLo, EnhPois, and EnhPr). We also observed a preference for temporal transitions into or out of the quiescent states (**Fig. 2b, c**).

To investigate whether the variations captured by the chromatin states could recapitulate the embryonic developmental trajectory, we applied the UMAP dimension-reduction technique^33^ to the 66 tissue biosamples using levels of chromatin marks at the genomic bins assigned to each chromatin state. H3K27ac signal levels at genomic bins assigned high-signal enhancers (Enh) in any of the 66 biosamples (in total 5.4% of the genome) cleanly segregated the 66 biosamples by tissue (**Fig. 2d, left panel**). The liver (with an endoderm origin) and heart (mesoderm) biosamples formed two separate clusters. Tissues with similar developmental origins were positioned near each other, with the four brain regions (ectoderm), the lung (endoderm) and the digestive organs stomach and intestine (endoderm), and limb and facial prominence (with cells from both endoderm and ectoderm origins) forming three clusters (**Fig. 2d**). The kidney (mesoderm) biosamples were positioned right next to the stomach, intestine, and lung (endoderm) biosamples. Furthermore, the earlier time-points (open symbols) are segregated from later time-points (filled symbols). A similar UMAP analysis on genomic bins assigned to the bivalent state (TssBiv) in any of the biosamples (in total 1.2% of the genome) by the levels of the ten chromatin marks also led to clear segregation of the biosamples by tissue, although there was some mixing between the lung biosamples and the stomach and intestine biosamples (**Fig. 2d, right panel**). Thus, the epigenomic landscapes captured by chromatin states Enh and TssBiv can accurately recapitulate the tissue lineages during embryonic development.

### Genome regions transit among TssBiv, Tss, and ReprPC states

Over developmental time, regions assigned to the bivalent promoter state (TssBiv), which has both active marks and the repressive H3K27me3 mark (**Fig. 1c**), can either lose repressive H3K27me3 and become active TSSs (Tss) or lose the active marks and transition into the repressive polycomb (ReprPC) state (**Supplementary Fig. 2**). Roughly 0.3% of any particular epigenome is assigned to the TssBiv state; cumulatively 1.2% of the genome is assigned to TssBiv across all tissues and time-points. TssBiv is less than half as prevalent as Tss and ReprPC, which constitute 0.8% and 0.8% of each epigenome and 2.2% and 5.5% of the genome overall, respectively. Almost all stretches of TssBiv genomic bins are flanked by ReprPC genomic bins. As an example, the promoter of the *Dlx1* gene is annotated as Tss in the forebrain, where it is highly expressed, and as TssBiv in the liver, where it is not expressed, and the bivalent promoter in the liver is surrounded by ReprPC regions (**Fig. 1e**). Among the genomic bins that are assigned TssBiv in any of the epigenomes, 64.7% are assigned ReprPC in at least one epigenome and 68.1% are assigned Tss in at least one epigenome (**Supplementary Fig. 3a**), indicating that a particular region is TssBiv in some tissue but becomes monovalent (Tss or ReprPC) in other tissues. Intriguingly, the overall fraction of TssBiv genomic bins decreased over the course of the development in all five tissues with seven time-points, although due to the small number of time-points this was statistically significant only in the three brain tissues (**Supplementary Fig. 3b**). This suggests that the resolution of TssBiv regions into a monovalent state is important for development, especially in the brain.

### Bivalent genes are involved in fundamental biological processes

We identified 14,558 bivalent regions, defined as stretches of TssBiv genomic bins surrounded by repressive chromatin states in any of the 66 biosamples (see **Methods**). These bivalent regions overlapped 14,729 GENCODE-annotated TSSs (**Supplementary Table 2**), belonging to 6,797 genes (**Supplementary Table 3**). There were 1,077 genes that were bivalent in all 12 tissues (i.e., having at least one bivalent TSS at one or more time-points of every tissue), and these genes were highly enriched in Gene Ontology (GO) terms related to embryonic development of myriad organs and systems, regulation of fundamental cellular processes, and modulation of cell-cell communications (**Supplementary Fig. 4a** and **Supplementary Table 4a, b**).

The liver had 5,482 bivalent genes (i.e., having at least one bivalent TSS at one or more time-points), 74% more than the other 11 tissues on average, and 1,291 of these 5,482 genes were not bivalent in the other 11 tissues. GO analysis on the 1,291 liver-only bivalent genes revealed terms that were involved in the development of a wide variety of organs other than the liver, such as heart, kidney, smooth muscle, brain, and cytoskeleton (**Supplementary Fig. 4b** and **Supplementary Table 4c, d**). We observed similar results for bivalent genes specific to other tissues. Thus, the bivalent genes in each fetal tissue reflect the regulatory pathways that are unused by the developmental program of that specific tissue.

### Bivalent genes exhibit repressed transcription

We further analyzed the expression of the 25,215 genes that were expressed (≥ 1 TPM) in at least one of the 66 biosamples, among which 6,324 were among our list of bivalent genes (**Methods**). We found that the bivalent genes in a tissue had lower expression levels than non-bivalent genes according to RNA-seq data in the same tissue. Across the 66 biosamples, the expression levels of bivalent genes were 5.2 ± 1.7 TPM, much lower than the expression levels of non-bivalent genes (39.8 ± 2.1 TPM; Wilcoxon rank-sum test P-value < 2.2×10^−16^). Furthermore, the genes that were not bivalent in any of the time-points of a tissue were expressed 7.79-fold higher (Wilcoxon rank-sum test P-values ≤ 2.2×10^−16^) than the genes that were bivalent at all time-points of the tissue (**Supplementary Fig. 5**). In a particular tissue, genes that were bivalent at different time-points were largely consistent (forebrain in **Fig. 3a**; all tissues in **Supplementary Fig. 6**). For example, 1,830 genes were bivalent at all seven time-points of the liver; only 439 such genes would be expected if the time-points were independent of one another (P-value < 2.2×10^−16^; Binomial test). Genes bivalent at the earliest time-point but not the latest time-point were expressed at significantly lower levels earlier in development; likewise, genes bivalent at the latest time-point but not at the earliest time-point were expressed at lower levels later in development (midbrain in **Fig. 3b**; all tissues in **Supplementary Fig. 7**). Both of these two sets of genes were expressed at significantly higher levels than genes bivalent at all time-points in the same tissue (**Fig. 3b**, **Supplementary Fig. 7**). Overall, the average expression level of a TSS across the time-points in a tissue is anti-correlated with the number of time-points at which the TSS is in a genomic bin assigned to the TssBiv chromatin state; in sharp contrast, a positive correlation is observed between expression and the duration the TSS is in a genomic bin assigned to the Tss chromatin state (**Fig. 3c**; **Supplementary Fig. 8**). Thus, the expression of bivalent genes is repressed in a tissue- and time-point-specific manner.

**Figure 3:**
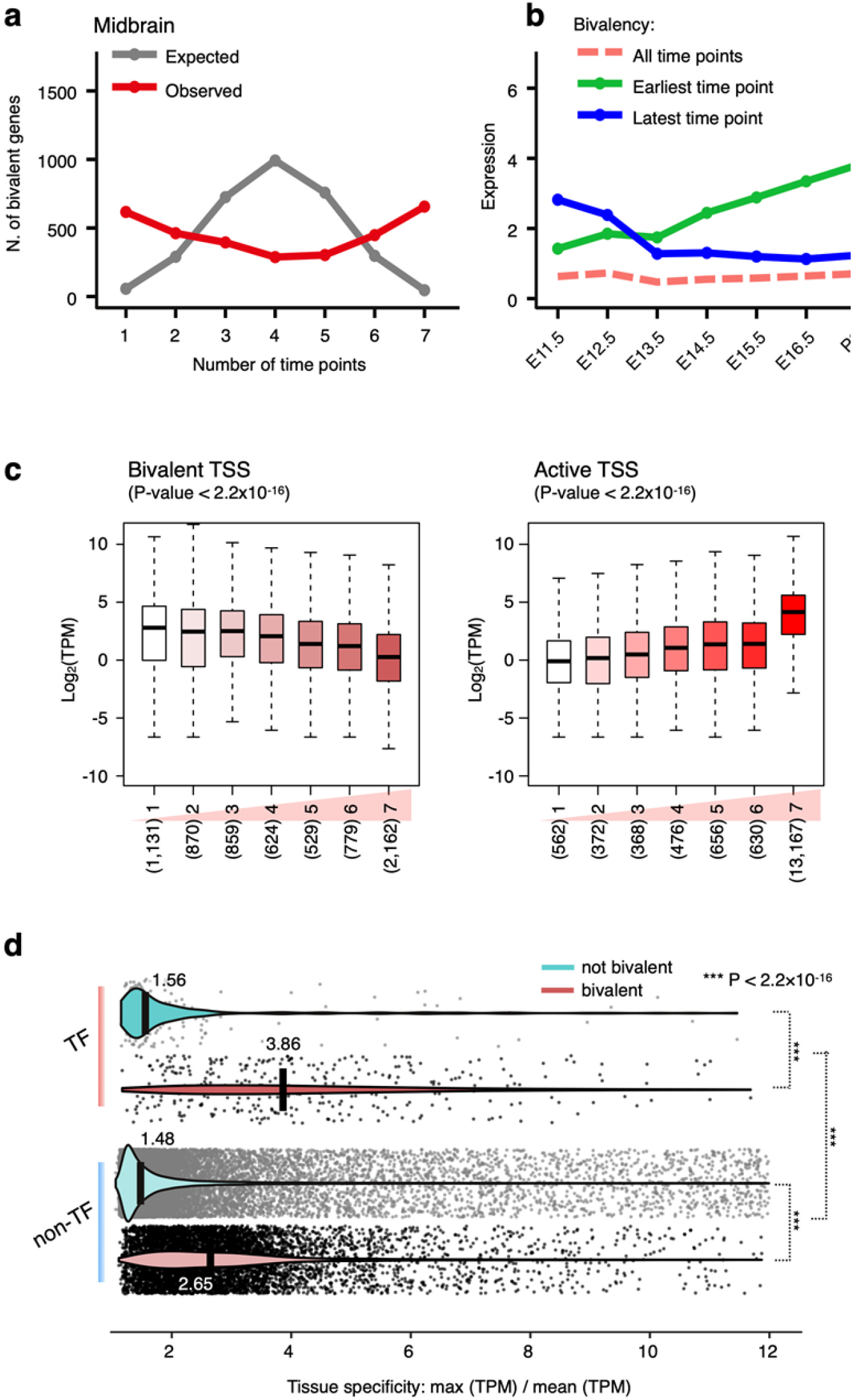
Count and expression of bivalent genes along developmental time-points. **a.** The number of bivalent genes at 1 to 7 time-points in the midbrain. Observed and expected numbers of genes are in red and in gray respectively. **b.** Median expression levels of three groups of genes: (green) bivalent at the earliest time-point but not at the last time-point, (blue) bivalent at the last time-point but not at the first time-point, and (pink) bivalent at all time-points. **c.** Distribution of gene expression, with genes grouped by the total number of time-points at which their TSSs are in the bivalent state TssBiv (left) or in the active state Tss (right) in the forebrain. For all box plots, whiskers show 95% confidence intervals, boxes represent the first and third quartiles, and the vertical midline is the median, and outliers are omitted. The total number of genes in each group is shown below each box plot in parentheses. There is a negative correlation between expression and the duration of the bivalent state and a positive correlation between expression and the duration of the active state (P-values < 2.2 × 10^−16^). **d.** Violin plots show the distributions of tissue-specificity scores for bivalent and non-bivalent genes that encode transcription factors (TFs) and non-TFs. Medians are shown in black bars with values indicated. P-values are shown for three comparisons as indicated.

### Bivalent genes are highly enriched in tissue-specific transcription factors

We compared the 6,797 bivalent genes (6,324 expressed in at least one of the 66 biosamples) with a curated list of 552 TFs with known DNA binding motifs in both mouse and human^34^, of which 535 were expressed in at least one of the 66 biosamples. A majority of the 535 TFs (338, 63.2%) were among the 6,324 bivalent genes (Chi-square P-value < 2.2×10^−16^). For both TF and non-TF genes, those that were bivalent were significantly more tissue-specific than those that were not bivalent (2.47-fold and 1.79-fold higher in median tissue specificity for TFs and non-TFs, respectively, Wilcoxon rank-sum test P-values < 2.2×10^−16^; **Fig. 3d**).

Consistent with earlier findings in embryonic stem cells^35,36^, a majority of the bivalent TSSs in our mouse fetal biosamples (mean = 62.5% across the 66 biosamples) overlapped CpG islands, much higher than non-bivalent TSSs (mean = 29.8%; Chi-square P-values in all 66 biosamples < 2.2×10^−16^). The enrichment is highly significant for the TSSs of both the TF genes (mean = 64.4% for bivalent TSSs vs. 43.5% for non-bivalent TSSs; P-values < 2.2×10^−16^) and the non-TF genes (62.3% vs. 29.5%, P-value < 2.2×10^−16^). CpG promoters are known to be less tissue-specific than non-CpG promoters^37^, which may seem at odds with our above finding that bivalent genes were significantly more tissue-specific than non-bivalent genes (**Fig. 3d**). To investigate the apparent contradiction, we separated bivalent and non-bivalent TSSs into CpG and non-CpG sub-groups. Indeed, each CpG sub-group is significantly less tissue-specific than the non-CpG subgroup with the same valency, yet the bivalent group is significantly more tissue-specific than the non-bivalent group when CpG and non-CpG promoters are combined (**Supplementary Fig. 9**).

We examined the TFs with the highest tissue-specificity scores, and a vast majority of these TFs were bivalent. Seventy-five TFs had tissue-specificity scores higher than 6, meaning that the highest expression level was at least as high as the expression levels in all other tissues combined (**Methods**). Of these, 64 were bivalent and the other 11 were not; we illustrate their tissue-specific gene expression (**Fig. 4a**) and the chromatin state assignments around eight example TFs (**Fig. 4b-i**). Two paralogous TFs, *Gata4,* and *Gata1* (**Fig. 4d, e**), illustrate bivalent and non-bivalent genes. *Gata4*, a bivalent gene, is predominantly expressed in the heart, consistent with its well-known role in regulating cardiac development^38^; it is also expressed at low levels in the stomach and intestine but not in other tissues. Accordingly, its TSS shows broad regions of the Tss state in the heart and narrower Tss regions surrounded by TssBiv and ReprPC regions in the stomach and intestine, while the TSS is covered by only TssBiv and ReprPC regions in other tissues (**Fig. 4e**). In comparison, *Gata1*, a non-bivalent gene, is a key regulator of erythrocyte development^39^ and is predominantly expressed in the liver. Consistently, the non-bivalent TSS of *Gata1* shows a broad Tss domain in the liver and a narrow Tss domain during early time-points of heart, but it is labeled Quies in other tissues (**Fig. 4d**). Thus, there are two distinct modes of gene repression: bivalent TSSs or quiescent TSSs.

**Figure 4:**
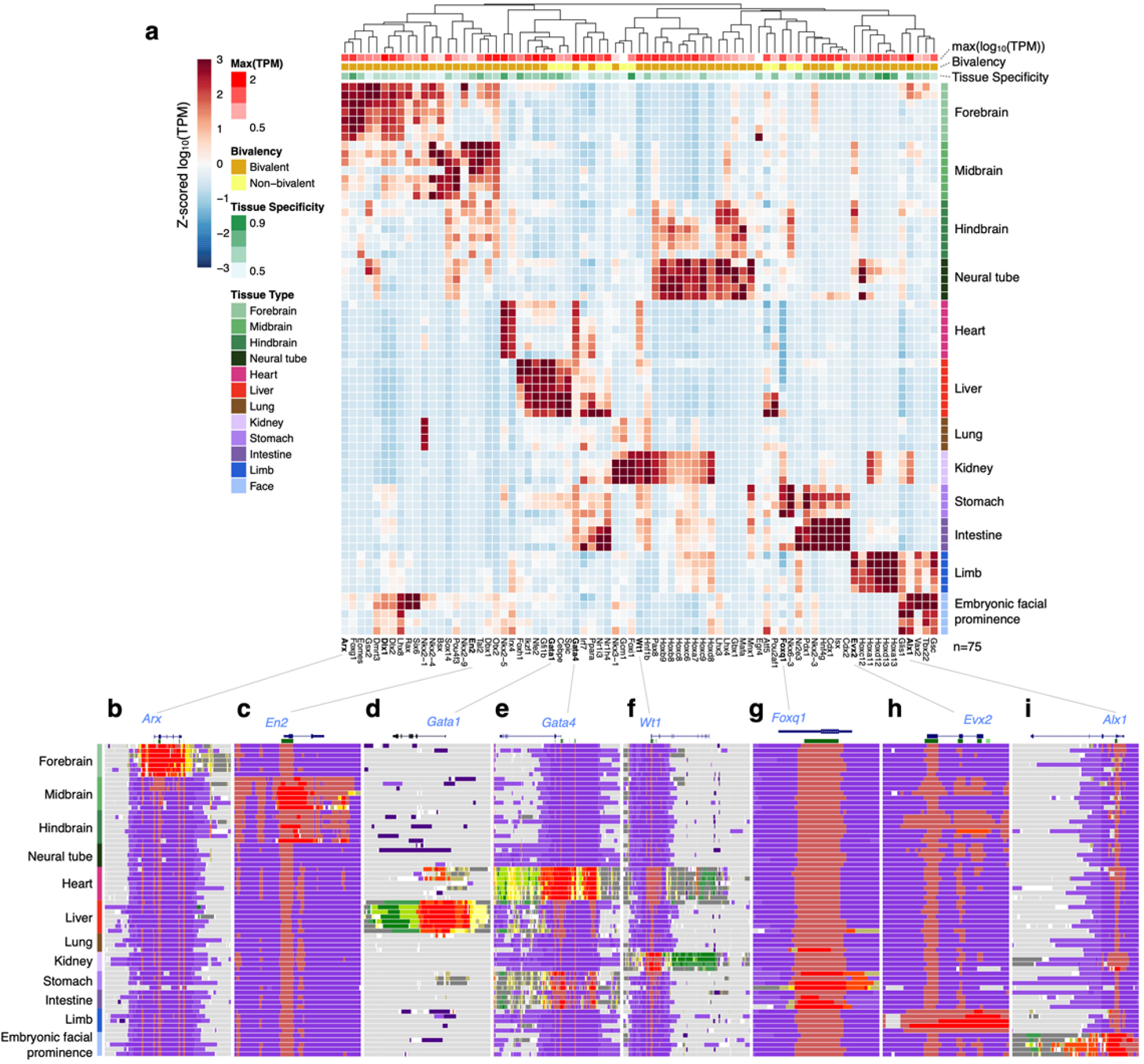
Expression profiles and chromatin states for the transcription factors with the highest tissue-specificity scores. **a.** Hierarchical clustering of expression profiles for the TFs with tissue specificity scores greater than 6, with 75 TFs in total. Rows on the top show the maximal expression level across all biosamples (intensities of red), bivalency status (brown for 62 bivalent TFs, and yellow for 13 non-bivalent TFs), and tissue specificity score (intensities of green). **b-i.** Example TFs and the chromatin state assignments near their loci. Among these, Gata1 (**d**) is a non-bivalent TF and the rest are bivalent TFs: Arx (**b**), En2 (**c**), Gata4 (**e**), Wt1 (**f**), Foxq1 (**g**), Evx2 (**h**), and Alx1 (**i**). Each gene name is near the 5′-end of the gene, and CpG islands are indicated as green boxes beneath each gene. Chromatin states are colored as in Fig. 1.

Other bivalent TFs show similar tissue specificity in their chromatin patterns—adopting the Tss state in the tissues where they are expressed while being in the TssBiv state flanked by ReprPC regions in the tissues that they are not expressed. The homeobox-containing transcription factor *Dlx1* is required for the migration of progenitor cells from the subcortical telencephalon to the neocortex as well as the differentiation of these progenitors into GABAergic neurons^40^. It is expressed in the forebrain and facial prominence; accordingly, its TSS adopts a highly active state in these tissues and the TssBiv-ReprPC repressive states in other tissues (**Fig. 1e**). *Arx* is another homeobox-containing transcription factor (**Fig. 4b**) important for the maturation and migration of GABAergic interneurons, and loss-of-function mutations of *ARX* cause lissencephaly (smooth brain) in humans^41^. *En2* encodes a homeobox transcription factor that is expressed at high levels in Purkinje cells and it functions as a transcriptional repressor for neurodevelopment, and *En2* mutant mice display defective cerebellar patterning and a reduction of Purkinje cell number^42^. *En2* is expressed only in the midbrain and hindbrain and shows the corresponding tissue-specific chromatin patterns (**Fig. 4c**). Wilms’ tumor-1 (*WT1*), which encodes a transcription factor and RNA-binding protein, is essential for kidney development^43^. It is predominantly expressed in the kidney and at lower levels in the heart, stomach, and intestine. Its TSS is in the Tss state in the kidney and shows a broad TssBiv domain in the heart while being TssBiv-ReprPC in other tissues (**Fig. 4f**). The forkhead transcription factor *Foxq1* is required for the maturation of the abundant mucin-producing foveolar cells that line the mucosal surface in the developing gastrointestinal tract^44^. *Foxq1* is expressed in the gastrointestinal tissues and in the Tss state in these tissues, but bivalent in other tissues (**Fig. 4g**). *Evx2* is required for the morphogenesis of limbs^45^, which is consistent with its expression and chromatin pattern (**Fig. 4h**). Finally, the aristaless-like homeobox 1 transcription factor *Alx1* plays an important role in the development of craniofacial mesenchyme, the first branchial arch, and the limb bud, and a complete loss of function of ALX1 protein causes severe disruption of early craniofacial development in humans^46^. Consistent with its functions, *Alx1* is predominantly expressed in the embryonic facial prominence and shows the corresponding chromatin profile (**Fig. 4i**).

### Genomic regions assigned to TssBiv are highly conserved evolutionarily

Genomic bins assigned to the bivalent state (TssBiv) are much more evolutionarily conserved than the genomic bins assigned to any of the other 17 chromatin states (**Fig. 5a**). In each biosample, we calculated the mean PhyloP^47^ score in each 200-bp genomic bin and then averaged these mean PhyloP scores for the genomic bins assigned to each chromatin state (**Methods**). The TssBiv state showed the highest PhyloP scores (0.51 averaged over the 66 biosamples), substantially higher (Wilcoxon signed-rank test P-values < 2.2×10^−16^) than the transcription-related states Tx (0.41) and EnhG (0.42), the active TSS state Tss (0.36), the high-signal enhancer state Enh (0.30), which were in turn substantially higher than the remaining 13 states, with Quies2 (0.02) being the lowest (**Fig. 5a**).

**Figure 5:**
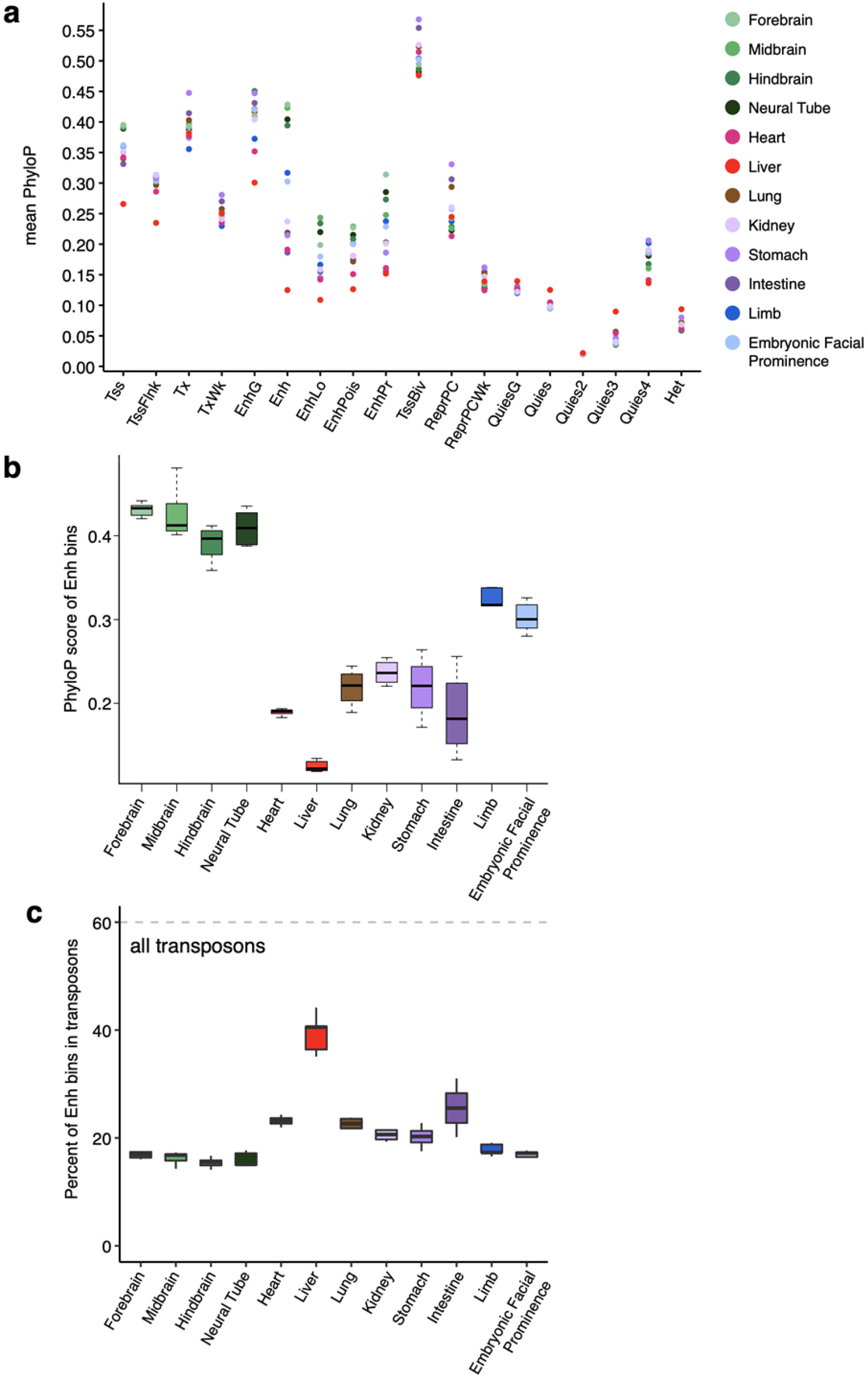
Evolutionary conservation of genomic regions by chromatin state. **a.** The PhyloP conservation score (phyloP60way for mm10) for genomic regions assigned to each chromatin state. Colors correspond to tissues. **b.** PhyloP score for genomic bins assigned to Enh in all 12 tissues. **c.** Percentage of bins assigned to Enh that overlap with transposons, for all 12 tissues.

For enhancer-related states (Enh, EnhLo, EnhPois, and EnhPr), the assigned regions in the four brain tissues (forebrain, midbrain, hindbrain, and neural tube) had the highest PhyloP scores, the regions in the liver had the lowest PhyloP scores, and the other seven tissues were in between (**Fig. 5a**). There were some variations in the PhyloP scores over the time-points within each tissue (**Supplementary Fig. 10**), but the four brain tissues were clearly the highest and the liver the lowest (**Fig. 5b**). For example, the average PhyloP score of Enh genomic bins was 0.42 for midbrain, while it was 0.13 for liver (Wilcoxon rank-sum test P-value = 5.8×10^−4^ for comparing the 7 midbrain time-points with the 7 liver time-points). We examined the transposon content in these Enh genomic bins and found that 40.6% of the Enh genomic bins in the liver overlapped annotated transposons, while only 14.1-17.5% of those in the four brain tissues did (**Fig. 5c**), which explained their substantially different levels of evolutionary conservation. These results suggest that the liver tissue has adopted some ancient transposon sequences as enhancers.

We directly examined the evolutionary conservation of the TSSs of TFs, stratified by whether they resided in a TssBiv genomic bin or not (the two bottom-right panels in **Supplementary Fig. 10**). The average PhyloP score of the TF TSSs in TssBiv genomic bins was 0.82, substantially higher than that of the TF TSSs not in TssBiv genomic bins (0.53, Wilcoxon rank-sum test P-value < 2.2×10^−16^ for comparing the two groups in 66 biosamples). Combined with our aforementioned findings that TFs are highly enriched in bivalent regions, these results indicate that TFs with bivalent TSSs play a key role in evolutionarily conserved pathways driving tissue development.

### Genomic regions assigned to TssBiv are enriched in PRC2-bound silencers and their target TSSs

We used a set of 1800 silencers bound by Polycomb Group 2 proteins (PRC2), identified using ChIA-PET assays targeting PRC2 component proteins in mouse embryonic stem cells^28^, to further annotate the chromatin states we defined in fetal mouse tissues. The PRC2-bound silencers overlapped extensively with the 14,558 bivalent regions (defined as TssBiv genomic bins surrounded by repressive bins; see **Methods**): 1069 out of 1800 silencers overlapped bivalent regions by at least 50% of the lengths of the silencers, while on average only 21 silencers overlapped with random regions with matching sizes as the bivalent regions (Z-score = 140; P-value < 2.2×10^−16^). In individual biosamples, the center locations of most silencers fall in the genomic bins assigned TssBiv or ReprPC (24 ± 4% and 28 ± 6% of the silencer centers, corresponding to 85.7- and 36.4-fold enrichment over the genomic footprints of these states), consistent with the enrichment of these two states in H3K27me3, the histone mark that PRC2 recognizes specifically.

The enrichment of the PRC2-bound silencers with chromatin states varied by silencer types. The silencers were clustered into four groups according to their H3K27ac signal profiles across the fetal mouse tissues^28^, a subset of the data we used to define chromatin states (H3K27ac is one of the ten marks used to train our ten-mark model). Group 1 silencers (N = 371) had the highest H3K27ac signals in the fetal mouse tissues^28^, and the centers of these silencers were in the Tss and Enh states in some biosamples, especially in the brain, but not so much in the liver (**Supplementary Fig. 11**). Group 2 silencers (N = 126) were depleted in H3K27ac in all fetal mouse tissues^28^, and the centers of most of these silencers were in quiescent states in all tissues (**Supplementary Fig. 11**). Group 3 and 4 silencers (N = 683 and 620) had intermediate levels of H3K27ac (higher in Group 3 than in Group 4)^28^, and their centers mostly fell in TssBiv and ReprPC states (**Supplementary Fig. 11**). We included in these alluvial plots the chromatin assignments in mouse embryonic stem cells (ES) using the ten-mark model with ENCODE data on 7 histone marks (missing H3K4me2, ATAC, and WGBS), which show similar chromatin state assignments as in the fetal tissues (**Supplementary Fig. 11**). To normalize for the genomic footprint of each genomic state, we compared genomic bins assigned to TssBiv (the least abundant state; **Fig. 1c**) with an equal number of genomic bins randomly drawn from the other states in individual biosamples for their overlap with each group of PRC2-bound silencers. TssBiv showed the highest enrichment for Group 1 and Group 3 silencers and moderate enrichment for Group 4 silencers; ReprPC showed moderate enrichment for all groups of silencers; Tss showed moderate enrichment for only Group 1 silencers; and none of the other states showed enrichment (**Fig. 6a**).

**Figure 6:**
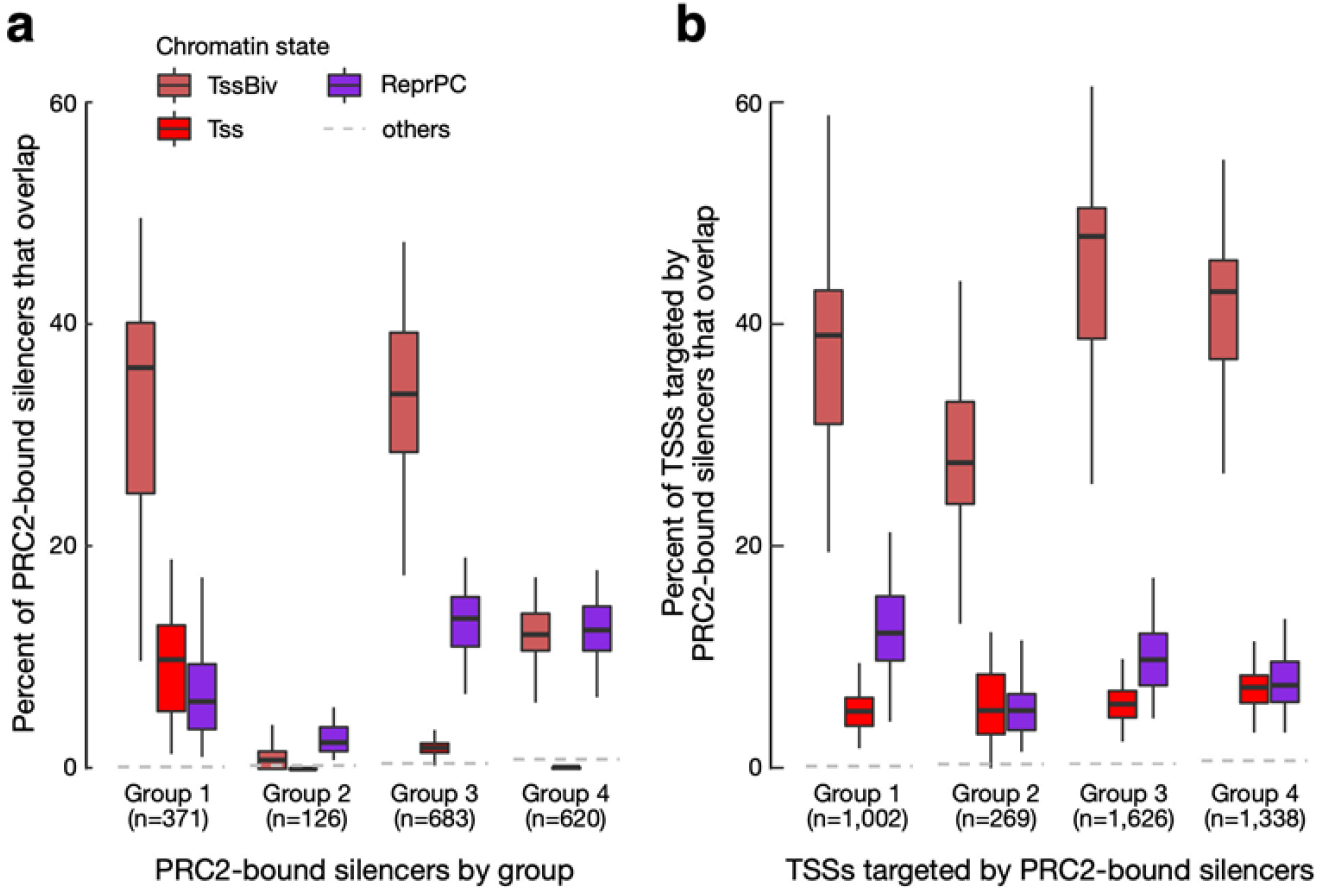
PRC2-bound silencers and their target TSSs are enriched in the TssBiv and ReprPC states. **a.** Percentage of PRC2-bound silencers whose centers overlap a genomic bin assigned to the TssBiv, Tss, ReprPC, or other chromatin states. Silencers were divided into four groups by Ngan et al.^26^ according to H3K27ac signals in mouse fetal tissue biosamples. To normalize for the differential genomic coverage of the chromatin states, the same numbers of genomic bins were randomly drawn in the other states to match the number of genomic bins in TssBiv in each biosample. States are colored as in Fig. 1b and the average of the other 15 states is shown as a gray dashed line. **b.** Same as **a** but for the TSSs targeted by the PRC2-bound silencers defined by Ngan et al.^26^.

The ChIA-PET data further provided the target TSSs for each PRC2-bound silencer^28^, and these TSSs also predominantly fell in the TssBiv, Tss, and ReprPC states, with the percentages of TSSs in active chromatin states ranked in the descending order for Group 1, 3, 4, and 2 silencers (**Supplementary Fig. 12**). Again, the brain regions showed higher percentages of TSSs in the Tss state than the liver for Group 1 silencers (e.g., 57.9% for forebrain and 13.4% for the liver; **Supplementary Fig. 12**). After normalizing for the genomic footprints of the chromatin states, TssBiv showed a strong enrichment for the target TSSs of all four groups of silencers, while Tss and ReprPC showed weak enrichment (**Fig. 6b**). Anong the 75 tissue-specific TFs (**Fig. 4a**), 44 of the 62 bivalent TFs but none of the 13 non-bivalent TFs were targeted by the PRC2-bound silencers (Fisher’s exact P-value = 1.6×10^−6^). Among the seven example bivalent TFs **(Fig. 4b-i**), five were targeted by the silencers (*En2*, *Gata4*, *Wt1*, *Foxq1*, and *Evx2*).

## DISCUSSION

We defined 18 chromatin states by integrating data on eight histone marks (ChIP-seq), chromatin accessibility (ATAC-seq), and DNA methylation (WGBS) in 66 biosamples across fetal mouse development (**Fig. 1**). We recapitulated the human states previously defined using fewer marks^10^ and refined enhancer, bivalent, and quiescent states. Regions annotated in these states showed higher variations among tissues and lower developmental variations across time-points in the same tissue (**Fig. 2a**), and the variations were specific enough to distinguish the tissue-of-origin for the 66 biosamples (**Fig. 2c**). Our chromatin state annotation should provide a useful resource for studying mammalian development.

We define two types of repressive states: ReprPC and ReprPCWk, the two states highly enriched in H3K27me3, jointly occupy 3.7% of the genome, and Het, the state highly enriched in H3K9me3, occupies 1.8% of the genome. However, Zaret and colleagues reported much larger genomic footprints for H3K27me3 domains (~10% of the human genome) and H3K9me3 domains (~20% of the human genome)^48^. They pointed out that if H3K9me3 and H3K27me3 ChIP-seq data were not normalized to input chromatin from the same experiment, reads for those marks would be under-represented, which could result in smaller H3K27me3 and H3K9me3 domains. We did normalize all histone mark ChIP-seq data to the input chromatin from the same experiment, and our signal files for H3K27me3 and H3K9me3 showed the same enriched regions as in the earlier work^48^; thus, the smaller genomic footprints of our ReprPC, ReprPCWk, and Het states were not a normalization artifact. We also define five quiescent states (Quies, Quies2, Quies3, Quies4, QuiesG) collectively occupy 80.5% of the mouse genome. These states show closed chromatin, very low levels of histone marks, and varying levels of DNA methylation. Except for Quies2, the other four quiescent states show low levels of H3K27me3 and H3K9me3 (**Fig. 1c**), the two repressive histone marks, and could encompass some of the H3K27me3 and H3K9me3 domains. Thus, we directly compared ChromHMM states with the H3K27me3 and H3K9me3 domains in the same IMR90 cell line as Becker et al., and found that 17% of H3K27me3 domains and 60% of H3K9me3 domains were in quiescent states; nevertheless, the ReprPC and ReprPCwk states were the most enriched in H3K27me3 domains and the Het state was the most enriched in H3K9me3 domain. Thus, Quies, Quies3, Quies4, and QuiesG states contain large portions of low-signal H3K27me3 and H3K9me3 domains.

Because enhancers and promoters have been examined extensively in previous ChromHMM studies^6,10,29^, we decided to focus on the TssBiv state in the current study. TssBiv has the smallest genomic footprint (0.3% of the genome in a particular biosample) among the 18 states, yet TssBiv is discovered consistently by the five-mark, eight-mark, and ten-mark models. TssBiv is particularly conserved evolutionarily, on average more conserved than genomic regions assigned to any other states (**Fig. 5**). We define 14,558 bivalent regions upon an integration of data in 66 biosamples, and roughly half of these regions overlap GENCODE-defined TSSs and the other half are intergenic. The bivalent TSSs show low mRNA levels in a tissue and developmental time-point specific manner (**Fig. 3**). These TSSs are highly enriched in tissue-specific TF genes (**Fig. 3, 4**). The TF TSSs in the TssBiv state are much more evolutionarily conserved than the TF TSSs in other chromatin states (the two bottom-right panels in **Supplementary Fig. 10**). Comparison with the recent ChIA-PET data^28^ revealed that the bivalent regions are highly enriched in PRC2-bound silencers and their target TSSs. Meanwhile, the TSSs of the target genes of PRC2-bound silencers are highly enriched in the TssBiv state in individual biosamples (**Fig. 6**). Taken together, these results indicate that TssBiv is a chromatin state that marks evolutionarily conserved PRC2-bound silencers and their target TSSs. It is intriguing that both PRC2-bound silencers and their target TSSs possess the same epigenetic signature and hence are assigned the same TssBiv state. This is perhaps not surprising because they are recognized by the PRC2 protein complex. Along this line of reasoning, Enhancers and active TSSs also share some epigenetic features (open chromatin, high levels of active marks such as histone acetylation, and low DNA methylation; Enh and Tss states in **Fig. 1c**).

Our systematic analysis of bivalent regions in mouse fetal tissues complement earlier studies on bivalent regions in other cell types and biological systems. Bivalent regions were first discovered in embryonic stem cells^23^, where their functions have been extensively studied. They have been shown to repress their associated genes and yet allow them to be poised for quick responses to stimuli. When embryonic stem cells differentiate, these bivalent genes become monovalent, retaining either the active marks or the repressive mark, and accordingly be expressed or repressed^19^. Subsequent studies reported bivalent domains in the differentiating CD4+ T cells^27^, the multipotent cranial neural crest cells^26^, adult intestinal villi cells with regenerative potential^24^, and terminally differentiated medium spiny neurons in the striatum^25^. In each of these studies, disruption of Polycomb group proteins led to the activation of the bivalent genes but not genes marked by H3K27me3 only^24,25^, suggesting that bivalency is a mechanism for persistent gene repression from embryonic stem cells to terminally differentiated cells.

Our analysis of bivalent genes in mouse fetal tissues indicates that they have low expression levels in the tissues where they are bivalent and are enriched for developmental transcription factors under tissue- and time-point-specific repression. A repressed gene can be in a quiescent chromatin state, which corresponds to low levels of all histone marks and high DNA methylation, such as GATA1 (**Fig. 4d**). Alternatively, it can be in an H3K9me3-enriched Het state accompanied by low levels of active histone marks and high levels of DNA methylation (**Fig 1d**). However, a majority of the bivalent TSSs in fetal tissues overlap CpG islands (mean = 62.5% across the 66 biosamples, vs. 29.8% for non-bivalent TSSs). DNA-hypomethylated CpG islands recruit both Polycomb group and Trithorax group proteins to lay down H3K27me3 and H3K4me3 marks respectively, and the expression level of the gene reflects the competition between Polycomb-mediated repression and Trithorax-mediated activation^49,50^. As a result, the interplay between the TssBiv, Tss, and ReprPC chromatin states (**Supplementary Fig. 3a**) reflects the main mechanism—distinct from quiescent or Het chromatin states—for silencing genes with CpG-rich TSSs in a tissue-specific manner throughout fetal development and possibly in adulthood.

In conclusion, we present genome-wide annotations of 18 chromatin states using ten chromatin marks all assayed in a mouse developmental matrix—twelve fetal tissues across 4-7 developmental time-points at daily intervals from E11.5 to birth. These comprehensive annotations enabled us to investigate the changes of chromatin profiles across tissue and time-points and connect the changes with gene expression. In particular, we analyzed bivalent regions in detail and found these evolutionarily conserved regions to be highly enriched in master transcriptional factors important for regulating tissue-specific developmental processes. More broadly, our results suggest that bivalent regions represent a mechanism for silencing CpG-rich genes in a tissue- and time-point-specific manner.

## METHODS

### Experimental data processing for mouse epigenome construction and chromatin state definition

We downloaded datasets processed for the mouse genome (mm10) from the ENCODE Portal^12,51^ (http://encodeproject.org) that corresponded to eight histone marks (H3K4me1, H3K4me2, H3K4me3, H3K9ac, H3K27ac, H3K36me3, H3K9me3, H3K27me3), ATAC-seq, and WGBS for each of 66 epigenomes (**Supplementary Table 1**). All biosamples were from the C57BL/6 mouse strain. For each histone mark, two biological replicates of the ChIP experiment were performed, and for each epigenome, two replicates of the control (input) experiment were performed. We ran ChromHMM^6^ on the 66 epigenomes at the default 200-bp resolution, using the histone ChIP-seq BAM files and the relevant control files for each dataset. For ATAC-seq data, each BAM file was converted to a signal track as follows. Reads were extended to their fragment size and counts-per-million were calculated for all non-overlapping 200-bp genomic windows. Quantile normalization was then applied across the entire data set and the normalized signal was binarized, using a threshold of 0.5. For WGBS data, BED files containing CpG percentages were downloaded from the ENCODE portal (**Supplementary table 1**), mean %CpG was calculated for all non-overlapping 200-bp genomic windows and after combining the two replicates for each biosample, binarization was applied, at a cutoff of 50% CpG.

We defined 18 chromatin states using ChromHMM^6^ using the processed data described above on the 10 marks and assigned each 200-bp genomic bin (13,627,678 of them in total for the entire mouse genome) to one of the 18 chromatin states in each biosample. We used the genomic bins with posterior probability > 0.5 for the downstream analysis; these bins composed 99% of the genome on average.

### Enrichment of chromatin states in other annotations (Fig. 1c)

We assessed the chromatin states assignments in each of the 66 epigenomes for their enrichments in three types of annotations (**Fig. 1c**, the right panel titled Enrichment): (1) for CpG islands, we downloaded cpgIslandExtUnmasked.txt from the UCSC Genome Browser; (2) we used GENCODE version M4 for gene-related annotations (transcription start sites or TSS, transcription end sites or TES, gene, exon, and intron); and (3) we used epigenetic annotations (EP300 and CTCF ChIP-seq peaks and DHS).

For every chromatin state, we computed its enrichment for each annotation, defined as the observed joint probability (P) of a chromatin state and an annotation occurring together over the expected joint probability (i.e., assuming the state and the annotation occur independently):

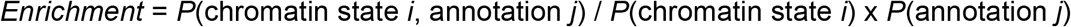

For visualization (the right panel of **Fig. 1c** titled Enrichment), the enrichments were scaled between 0 and 1:

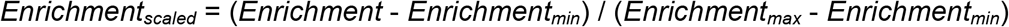

We further integrated the RNA-seq data (**Supplementary Table 1**) processed with the ENCODE uniform processing pipeline to compute the enrichment of the chromatin states in expressed or repressed genes for each of the 66 epigenomes^12^. For plotting the enrichment panels in **Fig. 1c**, we clustered genes into either expressed or repressed groups in each biosample based on an expression-level cutoff determined using a two-component Gaussian mixture model. The expression levels (in TPM) for the two replicates of each biosample were averaged.

We calculated the enrichment of the chromatin states in EP300 and CTCF ChIP-seq peaks and DNase hypersensitive sites (the right-most panel in **Fig. 1c**) for those epigenomes that had the EP300 and CTCF ChIP-seq or DNase-seq data available in the corresponding tissues and time-points (**Supplementary Table 1**). For the EP300 ChIP-seq data, the BAM files from two biological replicates were pooled, and peaks were called using MACS2^52^ with the q-value cutoff of 0.01. For the CTCF ChIP-seq data, the optimal IDR thresholded peaks^53^ defined by the ENCODE uniform ChIP-seq pipeline were used^12^. For the DNase-seq data, the hotspots defined by the ENCODE uniform DNase-seq processing pipeline were used^12^.

### Partial epigenome simulation and construction (Fig. 1d)

To assess the reliability of chromatin state assignments on epigenomes that lacked the data for one of the ten chromatin marks, for each biosample we simulated ten partial epigenomes, starting with the ten-mark epigenome and omitting the data for each mark individually. We applied the ten-mark 18-state ChromHMM model to the available data on the remaining nine marks and compared the resulting chromatin states assignments with the chromatin state assignments of the ten-mark epigenome by computing the Jaccard similarity between all genomic bins (**Fig. 1d**). The chromatin states with Jaccard similarity less than 0.5 were labeled as misassigned in the missing-one-mark epigenomes.

For the comparison with PRC2-bound silencers in embryonic stem cells, we also performed chromatin state assignment on embryonic stem cells, with data on seven histone marks (**Supplementary Table 1**), missing H3K4me2, ATAC, and DNA methylation data. We simulated the effect of missing three marks using midbrain and forebrain samples. These chromatin state assignments of the seven-mark epigenomes were used to define bivalent genes and compared with the bivalent genes defined using the chromatin state assignments of the ten-mark epigenomes (see below).

### Chromatin state variations across tissues and time-points (Fig. 2a)

We computed Jaccard similarity between a pair of epigenomes by comparing the chromatin states at the corresponding genomic bins between the two epigenomes.

### UMAP analysis of the epigenomes (Fig. 2d)

We performed two-dimensional visualization of the 66 epigenomes using UMAP^33^ analysis on two sets of 200-bp genomic bins: those assigned to the Enh state or the TssBiv state in one or more biosamples. For the Enh genomic bins, UMAP was provided with the H3K27ac signal levels across the 66 biosamples and the following parameters were used: n_neighbors = 7, min_dist = 0.5, seed = 11. For the TssBiv genomic bins, UMAP was provided with the signal levels of all ten marks across the 66 biosamples and the following parameters were used: n_neighbors = 10, min_dist = 0.04, seed = 12.

### Identification of bivalent TSSs and bivalent genes (Fig. 3, 4)

We developed a method to identify bivalent TSSs and bivalent genes by their chromatin states in each epigenome, described as follows. We first converted each epigenome to a character string using an 18-letter alphabet (one symbol for each state). Regular expressions were then used to extract punctate (median length 1800 bp) bivalent domains (stretches of contiguous genomic bins) in each epigenome, defined as bivalent chromatin states flanked by quiescent or heterochromatin states (ReprPC, ReprPCWk, Quies, Quies2, Quies3, Quies4, or QuiesG state). We used the union (14,558 regions across all tissue time-points, median 3,514 per tissue time-point, neighboring regions were not merged) of the detected genomic regions matching our regular expression for downstream analyses. Of the 14,558 regions detected in the 66 biosamples collectively, 14,729 regions overlapped GENCODE-annotated TSSs; we denote these *bivalent TSSs*. We further define a *bivalent gene* as having at least one bivalent TSS, yielding 6,797 genes that are bivalent in any of the 12 tissues.

We detected on average ~3,400 bivalent genes per tissue, defined as genes that are bivalent in any of the time-points in the tissue. We performed Gene Ontology (GO) analysis on bivalent genes using the PANTHER tool^54^. The genes used in the Gene Ontology (GO) analysis, of which the results are listed in **Supplementary Table 4** were obtained as follows: TSSs extracted from the M4 GENCODE annotations were intersected with the bivalent regions detected in each tissue. For each tissue, genes for which one or more TSSs intersected were retained. Then, the 1,077 genes that were found to have TSSs overlapping bivalent regions in *all* tissues were used as input for the GO analysis (**Supplementary Table 4a, b**). Another set of 1,291 genes was obtained using the same process, except genes were collected that had TSSs in bivalent regions *only* in liver samples and *not* in any other 11 tissues (**Supplementary table 4c, d**). Gene IDs were translated into gene names prior to submission to PANTHER. For six gene IDs, no matching gene name was found, leaving 1,074 and 1,288 genes in the “all tissues” and the “liver-only” gene sets for submission. PANTHER was run on the GO “Biological Process” ontology, using Fisher’s exact test and FDR for P-value calculations.

### Gene annotations and identification of transcription factors (Fig. 3d, Fig. 4, supplementary tables 2-4)

GENCODE M4 gene annotations were used to identify genes and transcription start sites (TSSs). To avoid double-counting TSSs, coinciding TSSs were merged. To identify transcription factors, we used the list of transcription factors and their homologs in mouse and human^34^. Ensembl IDs were obtained by mapping gene names to the GENCODE M4 annotations^55^. 552 TFs matched IDs in the GENCODE M4 mouse annotations.

### Evolutionary analysis (Fig. 5a-b, Supplementary Fig. 10)

We averaged the mouse 60-way phyloP^47^ score across the genomic positions in each 200-bp genomic bin. We then average this per-bin score for all the genomic bins assigned to a particular chromatin state in each biosample to obtain the average PhyloP score per state per biosample (**Supplementary Fig. 10**, first 18 panels). For each tissue (**Fig. 5a**), the PhyloP scores from the biosamples at different time-points were further averaged. For the TF TSSs (**Supplementary Fig. 10**, the two bottom-right panels), we used the PhyloP score for genomic bins where each TF TSS resided in, stratified by whether that bin was assigned to the TssBiv state or not.

### Overlap of Enh regions with annotated transposons (Fig. 5c)

We used transposon annotations in the mouse genome from Repbase^56^ to analyze the Enh state across different tissues (**Fig. 5c**). We overlapped the genomic bins assigned to the Enh state in each biosample with annotated transposons, requiring at least 1-bp overlap. The percentage of all genomic bins that overlapped transposons was used as control (gray dashed line in **Fig. 5c**).

### Analysis of PRC2-bound silencers (Fig. 6, Supplementary Fig. 11, 12)

We used the 18,000 PRC2-bound silencers classified into four groups based on their H3K27ac signal in mouse fetal tissues^28^. We overlapped the PRC2-bound silencers with our 14,558 bivalent regions, requiring at least half of the length of a silencer length to overlap. We randomly selected genomic regions with the same lengths as the bivalent regions to act as controls. Furthermore, we assigned each silencer to a chromatin state in a particular biosample according to which chromatin state the center of the silencer falls in.

We included embryonic stem cells in this analysis (ES-Bruce4). These cells were derived from C57BL/6, the same strain of mice from which the tissues were harvested. We only had data on seven histone marks on embryonic stem cells (**Supplementary Table 1**), and simulation of this partial epigenome (see above Methods) showed no major impact on the assignment of the TssBiv state and the resulting bivalent genes. Simulating using midbrain and forebrain samples, we found that most bivalent genes were identified using the partial epigenome. For example, among the 2,250 bivalent genes in the midbrain E11.5 sample, 2,014 (89.5%) were identified using the partial epigenome.

## Data availability

All experimental data used in this paper can be accessed at the encode Portal (http://www.encodeproject.org/), using the accession IDs listed in **Supplementary Table 1**.

## Code Availability

The code used to extract genomic regions based on regular expression can be found on GitHub, at https://github.com/weng-lab/stateregexp.git.

## Data visualization via a UCSC track hub

We made a track hub (https://users.wenglab.org/vanderva/trackhub/chromhmmpaper/hub_0.txt) for the UCSC genome browser^57^ to visualize all the data and annotations used in this study listed below. The trackhub can be accessed via a UCSC session: https://genome.ucsc.edu/s/Kaili/ChromHMM_paper.

1. ten-mark, 18-state chromatin state assignments (in dense mode) BigWig experimental data complete for 66 biosamples (in hide mode):

a. ChIP-seq of eight histone marks
b. ATAC-seq
c. WGBS
d. RNA-seq
e. DNase when available
f. EP300 ChIP-seq when available
g. CTCF ChIP-seq when available
2. ES-Bruce4 chromatin state assignments (in dense mode) BigWig experimental data for ES-Bruce4 (in hide mode)

a. ChIP-seq of seven histone marks
b. RNA-seq
c. EP300 ChIP-seq
d. CTCF ChIP-seq
3. Turn on the GENCODE gene annotation (in pack mode)
4. Turn on the CpG island track from UCSC (in dense mode)
5. Bivalent regions (in dense mode)
6. PRC-bound silencers and their target TSSs in two tracks (in dense mode)
7. Turn on the PhyloP conservation track (in full mode)
8. Turn on VISTA enhancer track hub (in hide mode)
9. Mouse cCREs (in hide mode)

## Acknowledgments

We thank ENCODE Consortium members for generating the ATAC-seq, ChIP-seq, RNA-seq, WGBS, and DNase-seq data on the 66 mouse embryonic biosamples and making them freely available. This work was supported in part by the National Institutes of Health grants HG009446 and HG007000 to ZW.

## Author contributions

AV: computational analysis, writing; KF: computational analysis; JT: computational analysis; JM: computational analysis; MP: computational analysis; HP: computational analysis, writing; ZW, project conception, design and management, writing.

## SUPPLEMENTARY FIGURE CAPTIONS

**Supplementary Fig. 1:**
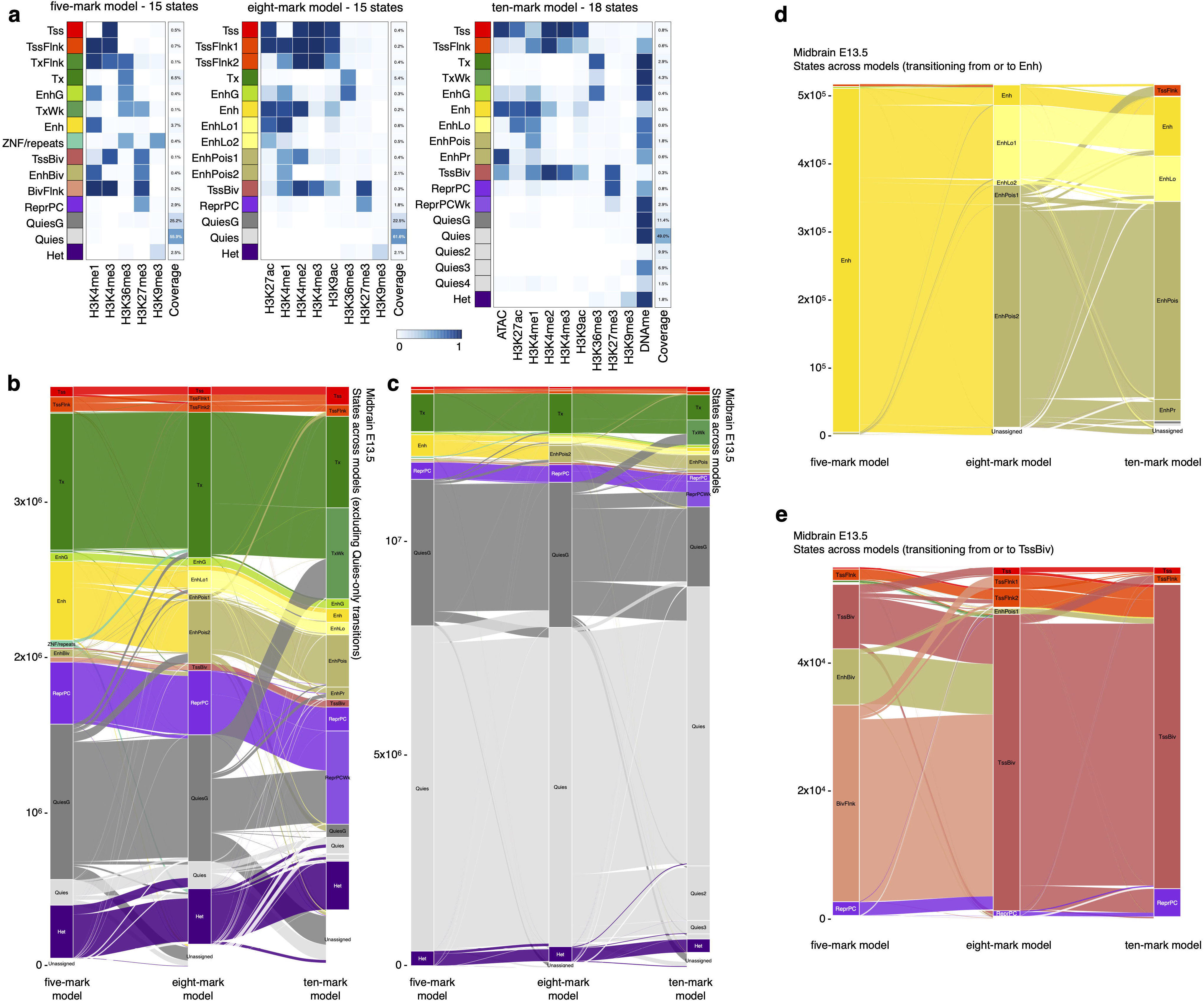
Comparison of the five-mark, eight-mark, and ten-mark models. **a** Emission probabilities for the five-mark 15-state model, the eight-mark 15-state model, and the ten-mark 18-state model, with the ten-mark model reproduced from **Fig. 1c** for easy comparison with the other two models. **b-e** Alluvial plots illustrate the correspondence of chromatin states across the three models in forebrain e13.5. **b** With genomic bins assigned to a quiescent state by all three models omitted, 3,743,342 genomic bins are shown. **c** All 13,627,678 200-bp bins in the genome. **d** Genomic bins assigned to Enh by one of the models. **e** Genomic bins assigned to TssBiv by one of the models.

**Supplementary Fig. 2:**
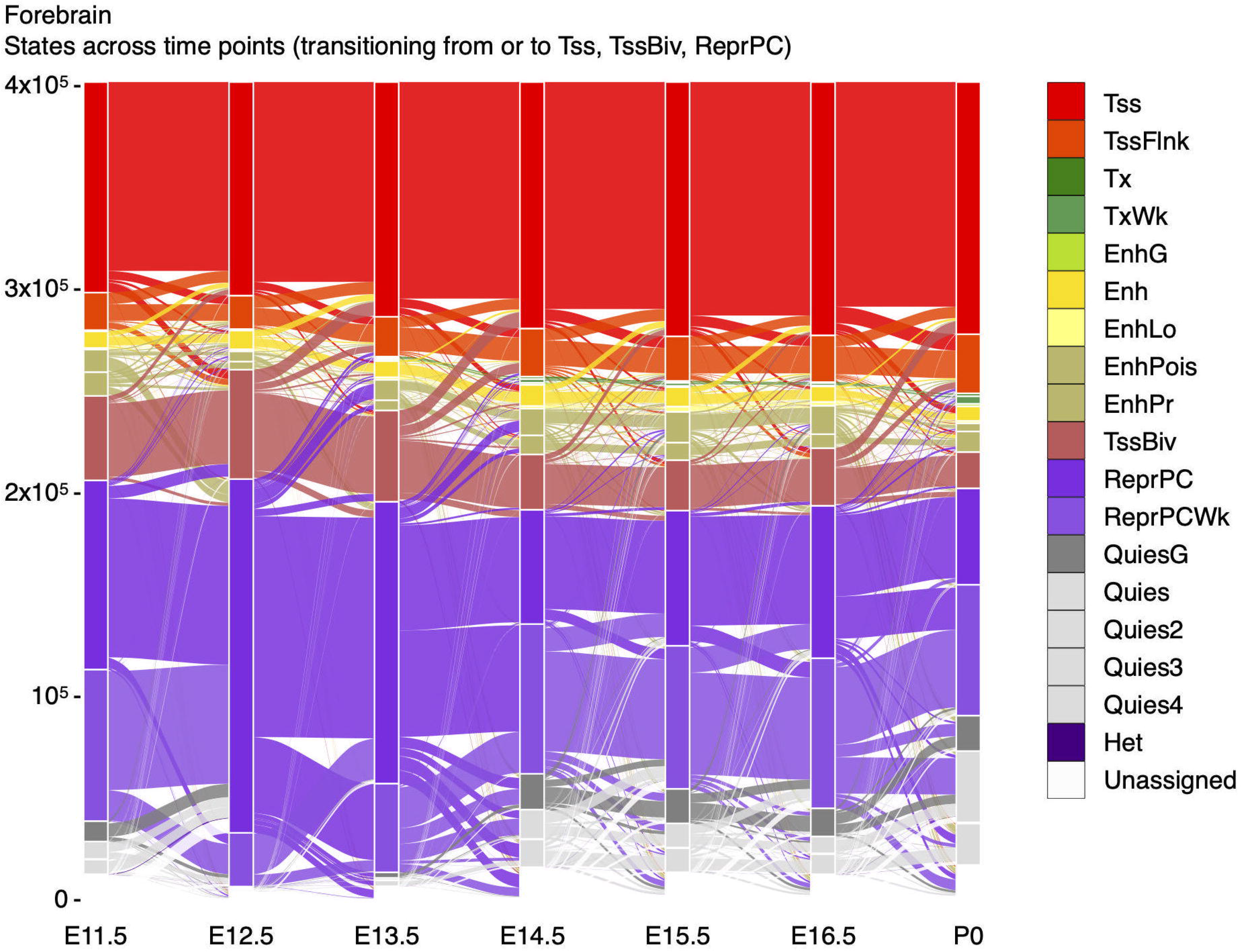
Chromatin state transition from early to late time-points. Genomic bins assigned to TssBiv in the forebrain at one or more time-points are included. States are colored as in Fig. 1c.

**Supplementary Fig. 3:**
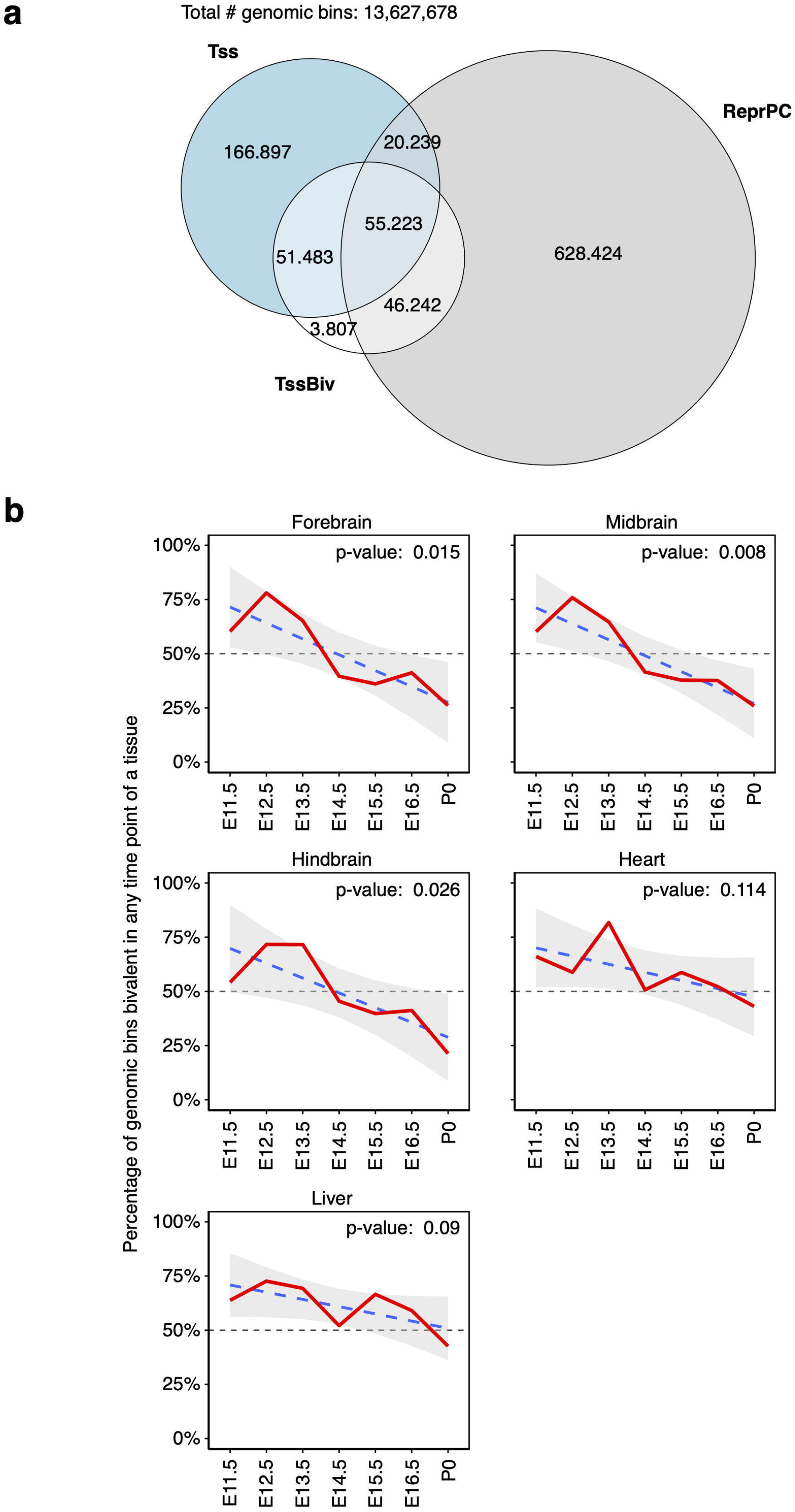
Comparison of genomic coverages of TssBiv, Tss and ReprPC, and the decrease of TssBiv coverage over time. **a** A Venn diagram shows the overlap of genomic regions assigned to the TssBiv, Tss, or ReprPC state in any of the 66 epigenomes. **b** red lines show the percentages of the genome in the TssBiv state over the time course of development for each tissue. Only the five tissues with seven time-points are included. P-values for linear fit (blue dashed line, with the 95% confidence interval in gray shaded area) are provided.

**Supplementary Fig. 4:**
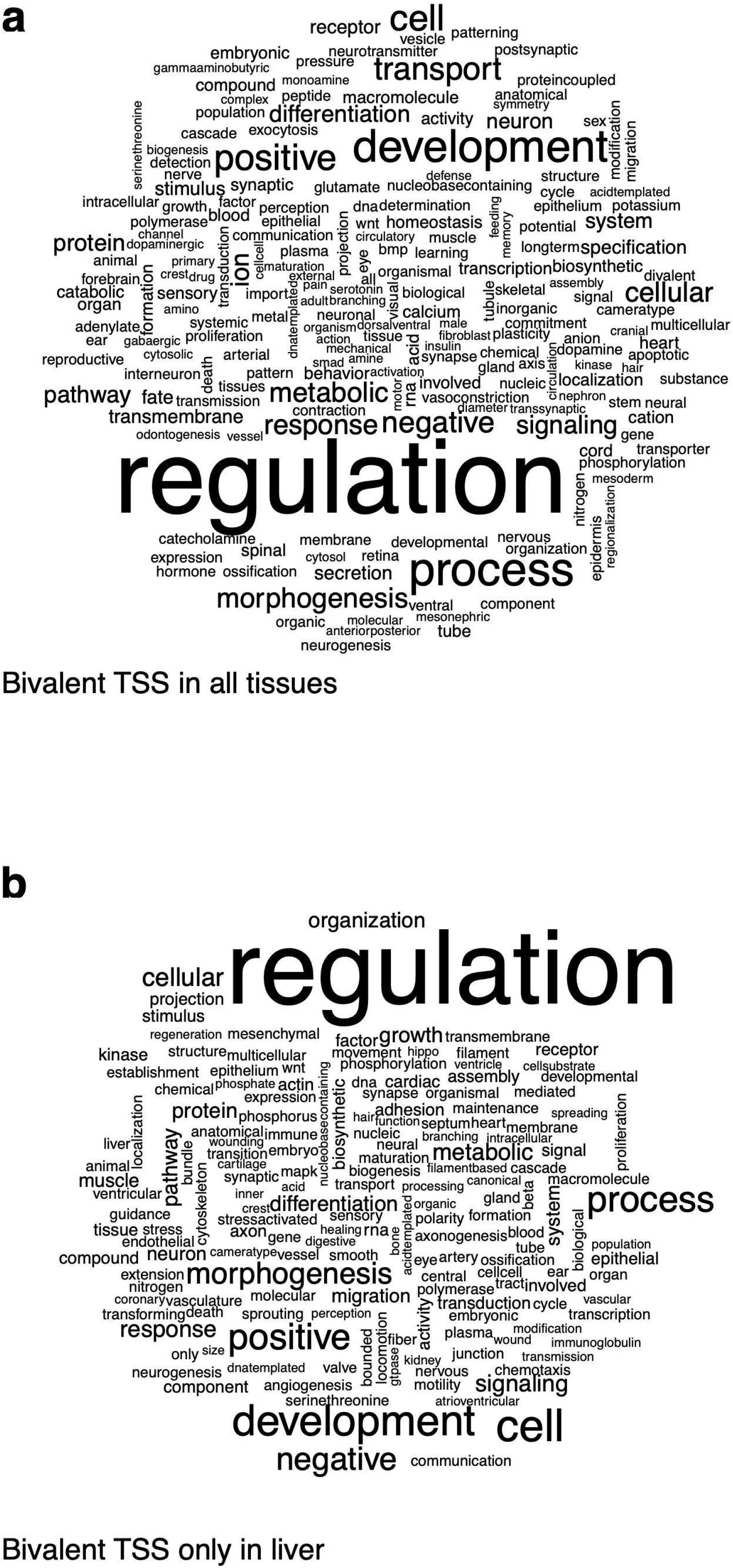
Word clouds for enriched GO terms in bivalent genes. Gene ontology (GO) enrichment analyses were performed using the PANTHER tool for two groups of genes: genes with bivalent TSSs in **(a)** all 12 tissues; **(b)** in the liver and not in any other tissues. For each analysis, a summary of significantly enriched GO terms is presented as a word cloud. See Supplementary Table 4 for full PANTHER results.

**Supplementary Fig. 5:**
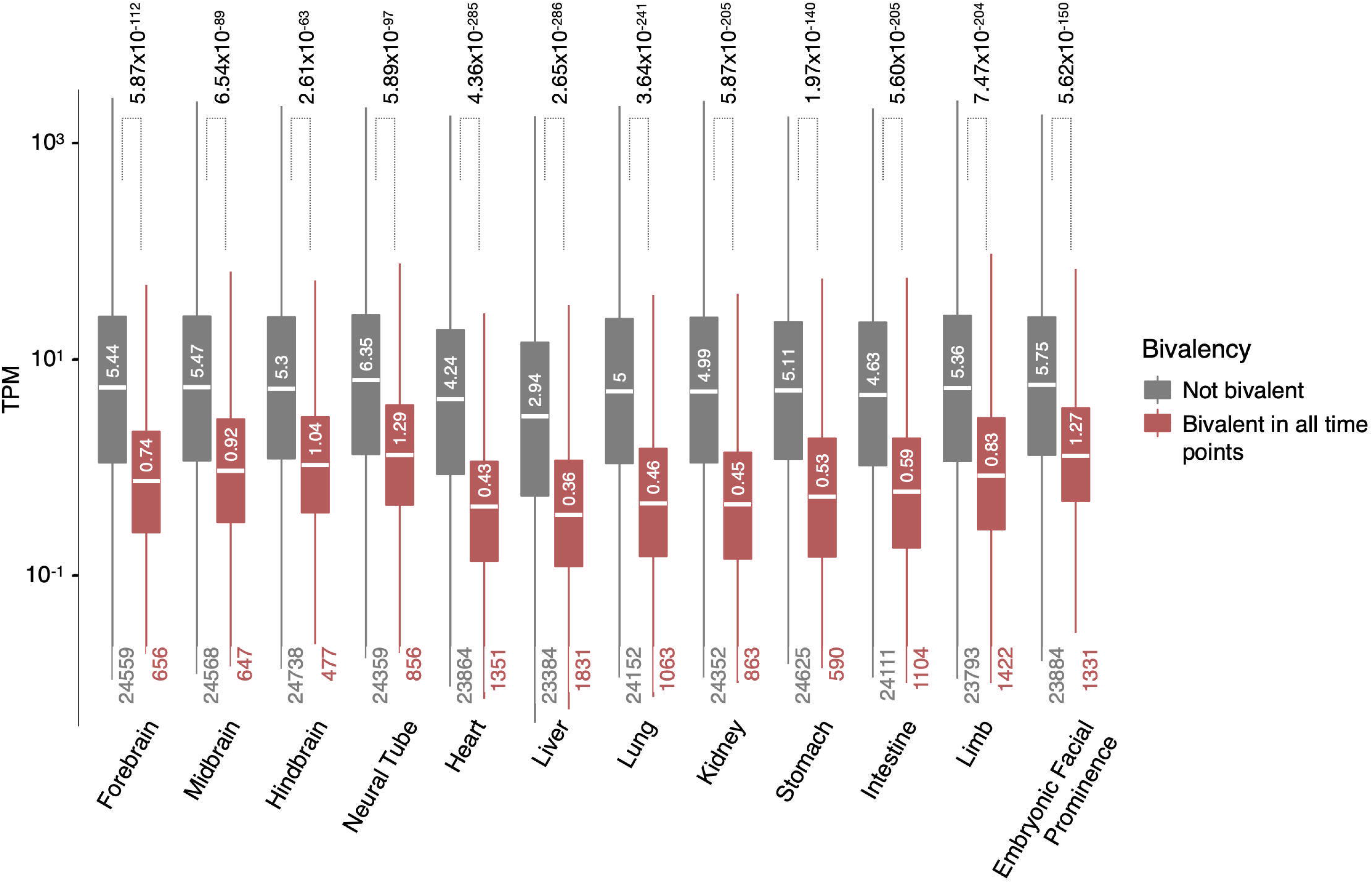
Expression levels of genes with or without bivalent TSSs. For each tissue, the expression levels of genes (in TPM) are plotted, stratified by whether it has a bivalent TSS at all time-points. For each box plot, the total number of genes in each group is shown at the bottom. Outliers are omitted for clarity. Wilcoxon P-values for comparing the two groups of genes in each tissue are provided.

**Supplementary Fig. 6:**
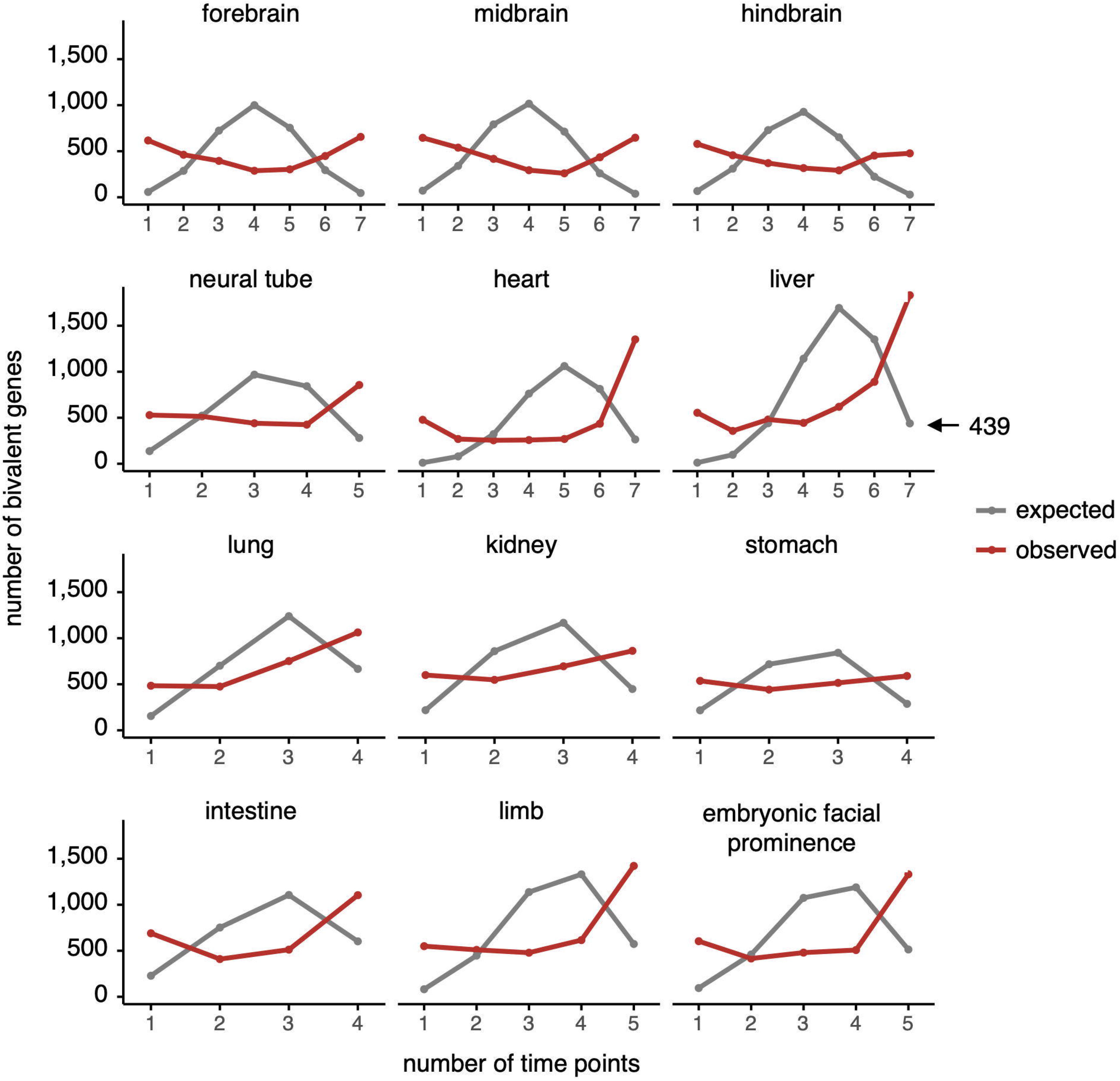
Number of genes that are bivalent in a certain number of time-points. For each tissue, the total number of genes that are deemed bivalent in a certain number of time-points is plotted in red, compared with the expected number (in grey) if genes were randomly assigned to be bivalent at each time-point.

**Supplementary Fig. 7:**
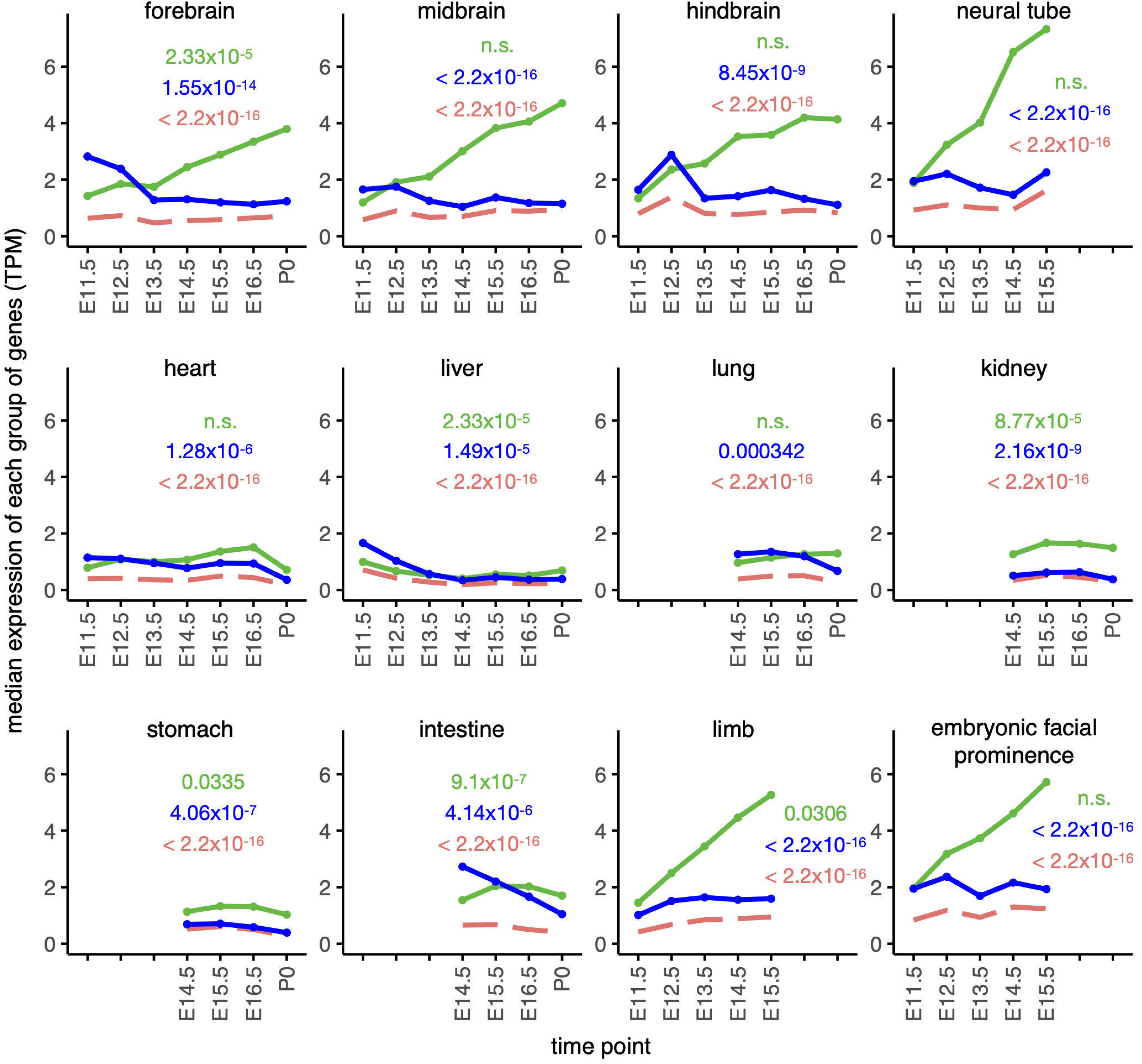
Expression of genes with bivalent TSS at early vs. late time-points. Median expression levels of genes stratified into three distinct categories are plotted: genes deemed bivalent at the first time-point but not at the last (early-bivalent genes; blue line); genes deemed bivalent at the last time-point but not at the first (late-bivalent genes; green); and genes with bivalent TSS at all time-points (all-bivalent genes; red dashed line). Wilcoxon rank-sum test P-values for the comparisons between early- and late-bivalent genes for their expression levels at the first time-point (green P-values); between the early- and late-bivalent genes at the last time-point (blue P-values); and between all-bivalent genes vs. early- and late-bivalent genes (red P-values). n.s. stands for not significant.

**Supplementary Fig. 8:**
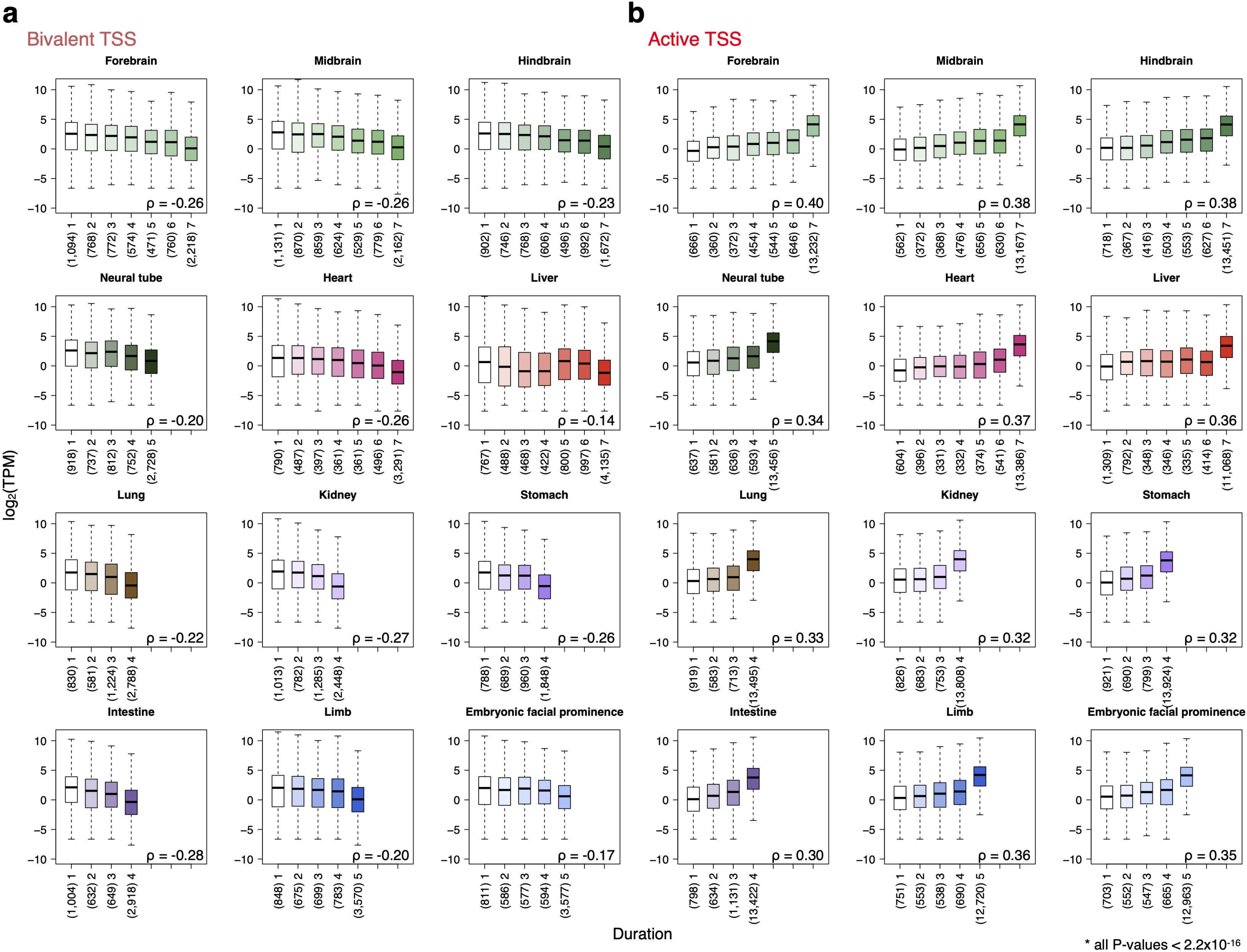
Correlations between gene expression and the duration of bivalent and active TSSs. The expression levels of genes with a certain duration (number of time-points) of being bivalent (**a**) or active (**b**) in each tissue. Numbers in parentheses indicate the number of genes for each duration. P-values were computed with ANOVA with multiple-testing correction.

**Supplementary Fig. 9:**
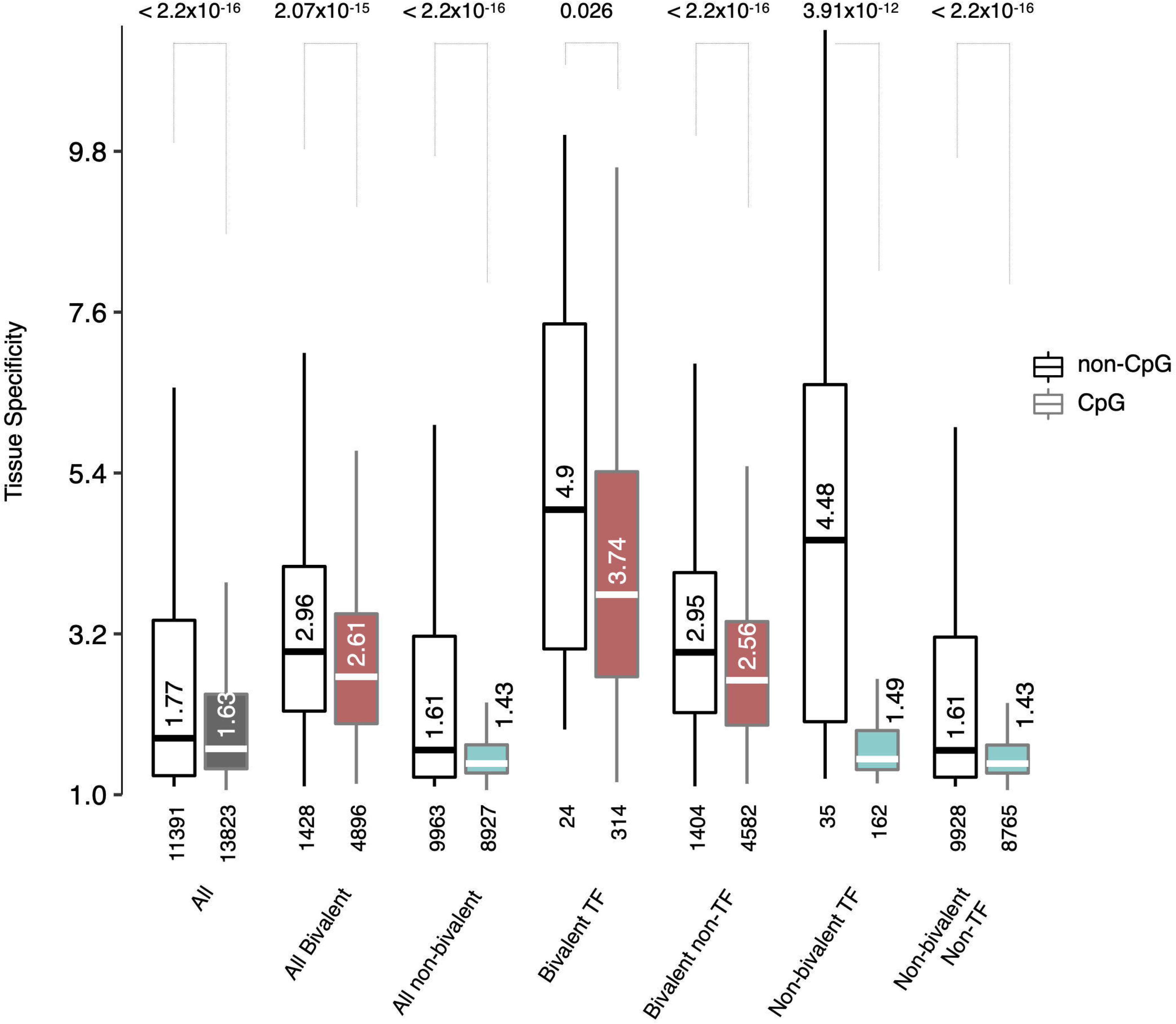
Tissue specificity for groups of genes classified by whether their TSSs overlap CpG islands. Tissue-specificity scores are shown for all genes, and subsets of genes depending on whether they encoded TFs, they have a bivalent TSS, and whether the TSSs overlap CpG islands. Wilcoxon P-values for comparing the CpG and non-CpG groups of genes are provided.

**Supplementary Fig. 10:**
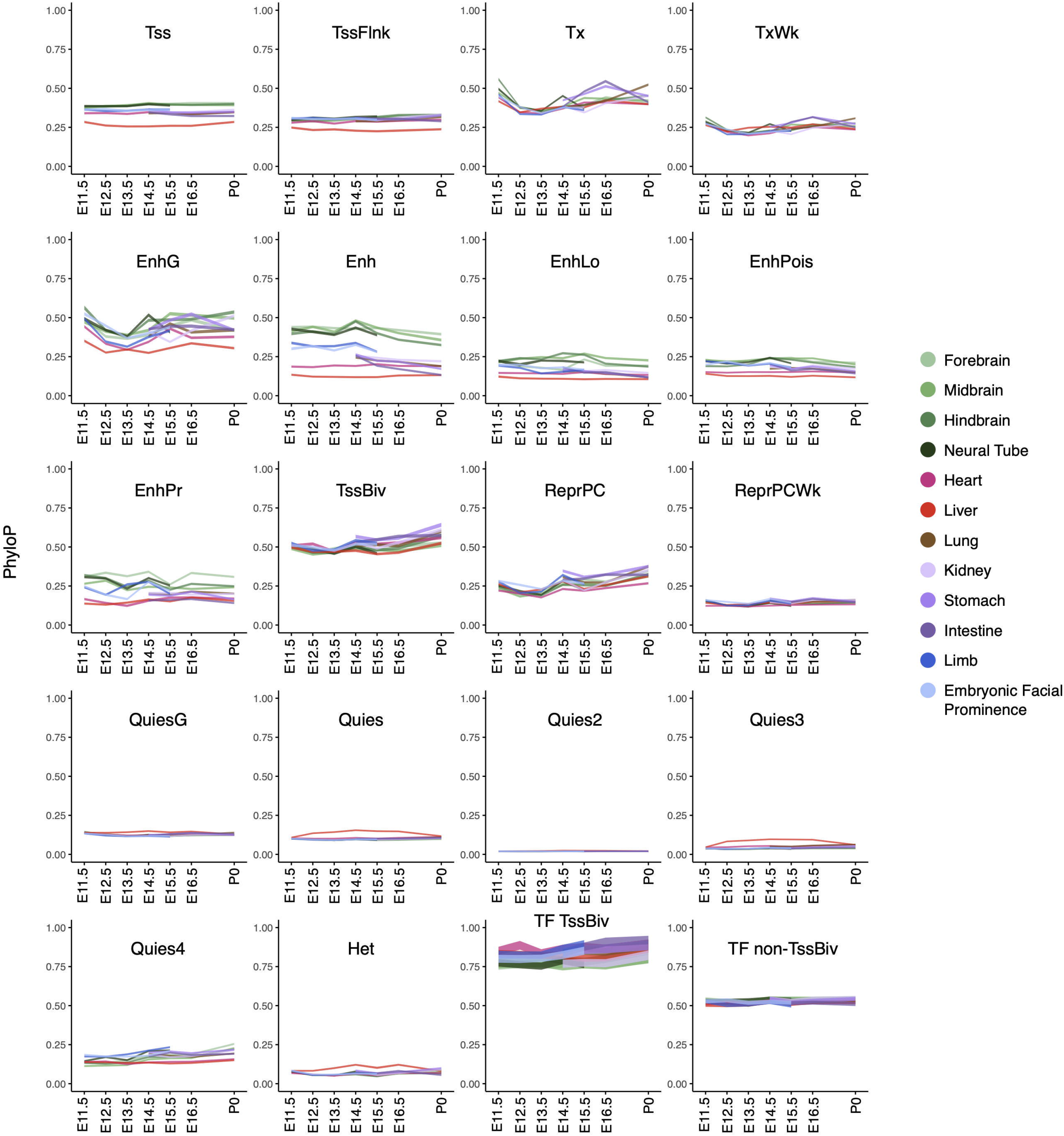
Evolutionary conservation for genomic bins assigned to each chromatin state in each biosample. Average PhyloP scores are plotted for the 18 chromatin states. The last two panels (bottom right) are TSSs of transcription factors stratified by whether they fall in a TssBiv genomic bin or not. The thickness of a line corresponds to the standard error. Tissues are colored accordingly.

**Supplementary Fig. 11:**
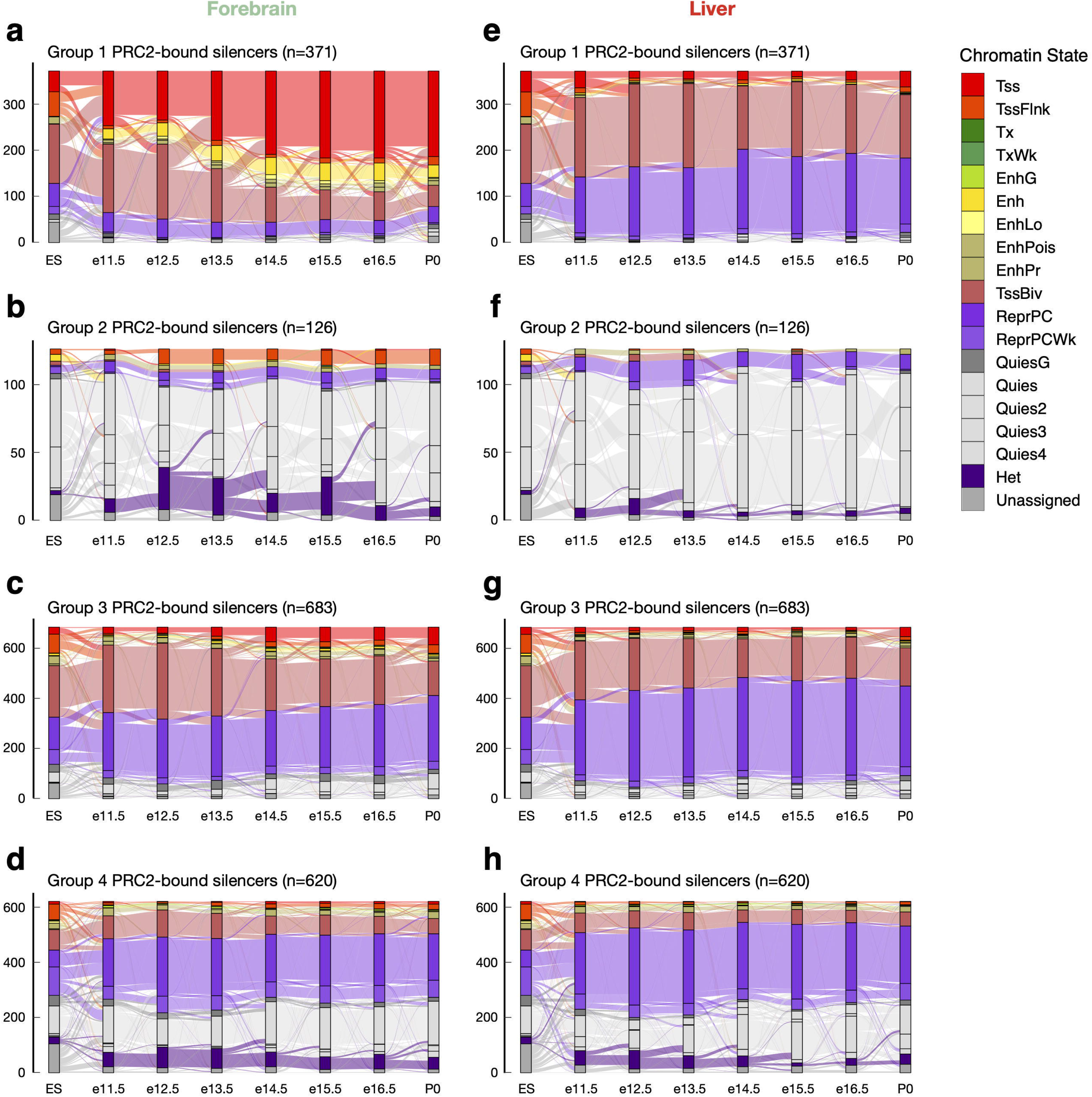
Chromatin state assignments for the center positions of PRC2-bound silencers. Four groups of PRC2-bound silencers correspond to those in Fig. 6a are plotted across time-points in the forebrain (**a-d**) and liver (**e-h**). The state assignments for mouse embryonic stem cells (ES) are included for comparison. States are colored as in Fig. 1c.

**Supplementary Fig. 12:**
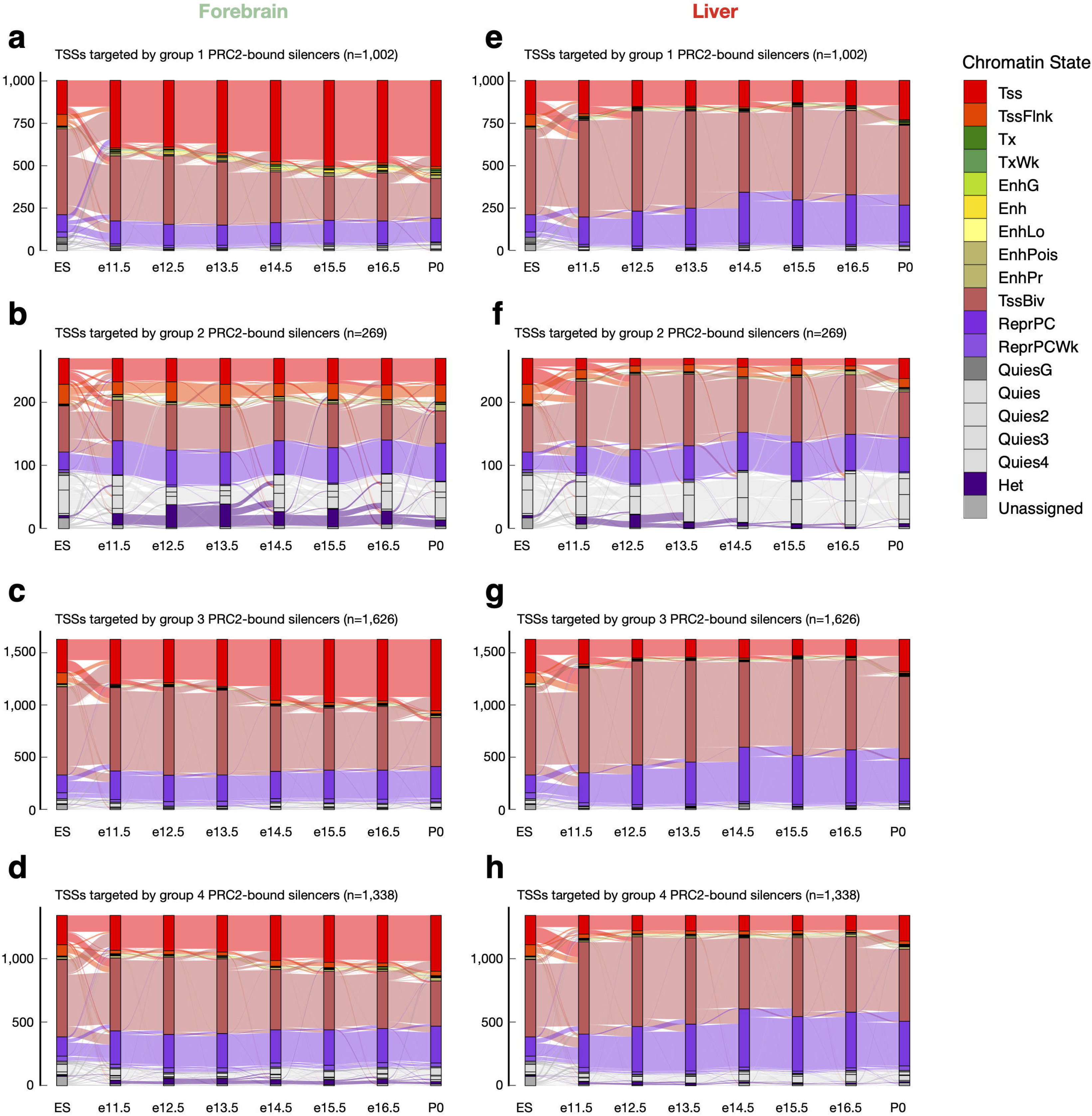
Chromatin state assignments for the TSSs targeted by PRC2-bound silencers. This figure corresponds to supplementary Fig. 12 but for the TSSs targeted by PRC2-bound silencers.

## SUPPLEMENTARY TABLES

**Supplementary Table 1: Input datasets and their ENCODE accessions.** ENCODE file accession IDs for all input files. **a.** BAM files for histone ChIP-seq datasets and controls. **b.** BED files with CpG calls from WGBS. **c.** RNA-seq TPM matrices for the two replicates of each biosample. **d.** BAM files for ATAC-seq. **e.** BAM files for DNase-seq.

**Supplementary Table 2: Bivalent TSSs in each biosample.** GENCODE M4 TSS annotations were intersected with bivalent regions in each biosample. Sites occupying the same genomic position were merged.

**Supplementary Table 3: Bivalent genes and their expression levels. a.** Expression levels are reported in TPM, in each tissue and time-point. **b.** The number of bivalent genes shared between any pair of biosamples. **c.** The number of bivalent genes shared between any pair of tissues. Diagonal numbers indicate the total number of bivalent genes in each tissue. **d.** Bivalent state of the TSSs of the genes in each biosample. **e.** Bivalent regions defined across all biosamples. **f.** Union of bivalent regions detected in all biosamples, as determined by regular expression (see Methods).

**Supplementary Table 4: GO enrichment analysis using the PANTHER tool. a.** PANTHER output for genes that are bivalent in all tissues. **b.** List of genes submitted for analysis in **a**. **c.** PANTHER output for genes that are bivalent exclusively in the liver. **d.**List of genes submitted for analysis in **c**.

